# Single-cell foundations of live social gaze interaction in the prefrontal cortex and amygdala

**DOI:** 10.1101/2021.08.25.457686

**Authors:** Olga Dal Monte, Siqi Fan, Nicholas A. Fagan, Cheng-Chi J. Chu, Michael B. Zhou, Philip T. Putnam, Amrita R. Nair, Steve W. C. Chang

## Abstract

Social gaze interaction powerfully shapes interpersonal communication in humans and other primates. However, little is known about the neural underpinnings of these social behavioral exchanges. Here, we studied neural responses associated with naturalistic, face-to-face, social gaze interactions between pairs of macaques. We examined spiking activity in a large number of neurons spanning four different brain regions involved in social behaviors – the amygdala, orbitofrontal cortex, anterior cingulate cortex, and dorsomedial prefrontal cortex. We observed widespread single-cell representations of social gaze interaction functionalities in these brain regions – social discriminability, social gaze monitoring, and mutual eye contact selectivity. Many of these neurons discriminated looking at social versus non-social stimuli with rich temporal heterogeneity, or parametrically tracked the gaze positions of oneself or the conspecific. Furthermore, many neurons displayed selectivity for mutual eye contact as a function of the initiator or follower of mutual gaze events. Crucially, a significant proportion of neurons coded for more than one of these three signatures of social gaze interaction, supporting the recruitment of partially overlapping neuronal ensembles. Our findings emphasize integrated contributions of the amygdala and prefrontal circuits within the social interaction networks in processing real-life social interactions.

## Main

Social behaviors involve both ‘social perception’, where each individual observes and gains information about others, and ‘social interaction’, where two or more individuals actively send and receive behavioral signals to and from one another over time (Argyle and Cook, 1976; Emery, 2000; Risko et al., 2016). In humans and non-human primates, gaze is an important behavioral modality underlying large parts of social perception and interaction. Social gaze interaction, powerfully guiding interpersonal communication (Emery, 2000; Itier and Batty, 2009; Kleinke, 1986), requires continuous gaze monitoring of both oneself and other (Hari et al., 2015; Redcay and Schilbach, 2019) in behaviorally contingent and communicative manners (Shepherd and Freiwald, 2018). However, while a great deal of knowledge exists regarding the circuitry underlying social gaze perception (Carlin et al., 2011; Haxby et al., 2000; Pelphrey et al., 2004; Perrett et al., 1985), we know very little about the neural mechanisms of social gaze interaction.

How the primate brain mediates social gaze interaction remains an open question. Interindividual gaze variables might be computed in a localized manner for efficient and modular processing with well-defined region-specific functionalities. Alternatively, these variables might be computed across multiple areas in the primate social interaction networks (Freiwald, 2020) for highly distributed and globally integrated processing. During naturalistic social gaze interactions in pairs of macaques, here we studied single-cell substrates of social gaze interaction in four distinct regions in the primate prefrontal and amygdala networks – basolateral amygdala (BLA), orbitofrontal cortex (OFC), anterior cingulate gyrus (ACCg), and dorsomedial prefrontal cortex (dmPFC). In the primate brain, BLA (Chang et al., 2015; Gothard et al., 2007; Grabenhorst et al., 2019; Mosher et al., 2014; Munuera et al., 2018; Pryluk et al., 2020), OFC (Azzi et al., 2012; Chang et al., 2013; Watson and Platt, 2012), ACC/ACCg (Basile et al., 2020; Chang et al., 2013; Haroush and Williams, 2015), and dmPFC (Falcone et al., 2017; Jamali et al., 2021; Noritake et al., 2018; Yoshida et al., 2012) have all been implicated in multitudes of functions associated social learning and decision-making. In these brain regions, here we focused on identifying single-neuron correlates of three key functionalities important for social gaze interaction, namely discriminating social from non-social stimuli, tracking one’s own and other’s gaze positions, and differentiating mutual from non-mutual gaze events.

### Social discriminability of prefrontal and amygdalar neurons during live social gaze interaction

Six pairs of rhesus macaques (M1: recorded monkey, ‘self’; M2: partner monkey, ‘other’) engaged in dyadic face-to-face social gaze interaction (Dal Monte et al., 2016, 2017) while the eye positions of both monkeys were simultaneously and continuously tracked at high temporal and spatial resolution (Fig. 1a, Fig. S1; Methods). We quantified spontaneously occurring naturalistic gaze behaviors in four gaze regions of interest (ROIs): *Face*, *Eyes*, *Non-eye Face* (i.e., face excluding the eye regions) and non-social *Object* (Fig. 1a–b). Replicating the significance of gaze directed to face and eyes in humans and non-human primates (Dal Monte et al., 2016; Gothard et al., 2004; Itier and Batty, 2009; Kano et al., 2018), the duration of each fixation was longer and the total number of fixations was higher when monkeys explored *Eyes* compared to *Non-eye Face* or *Object* (all p < 0.0001, Wilcoxon sign rank, two-sided, FDR-corrected) and when they explored *Non-eye Face* compared to *Object* (both p < 0.0001) (Fig. 1c).

**Figure 1.**
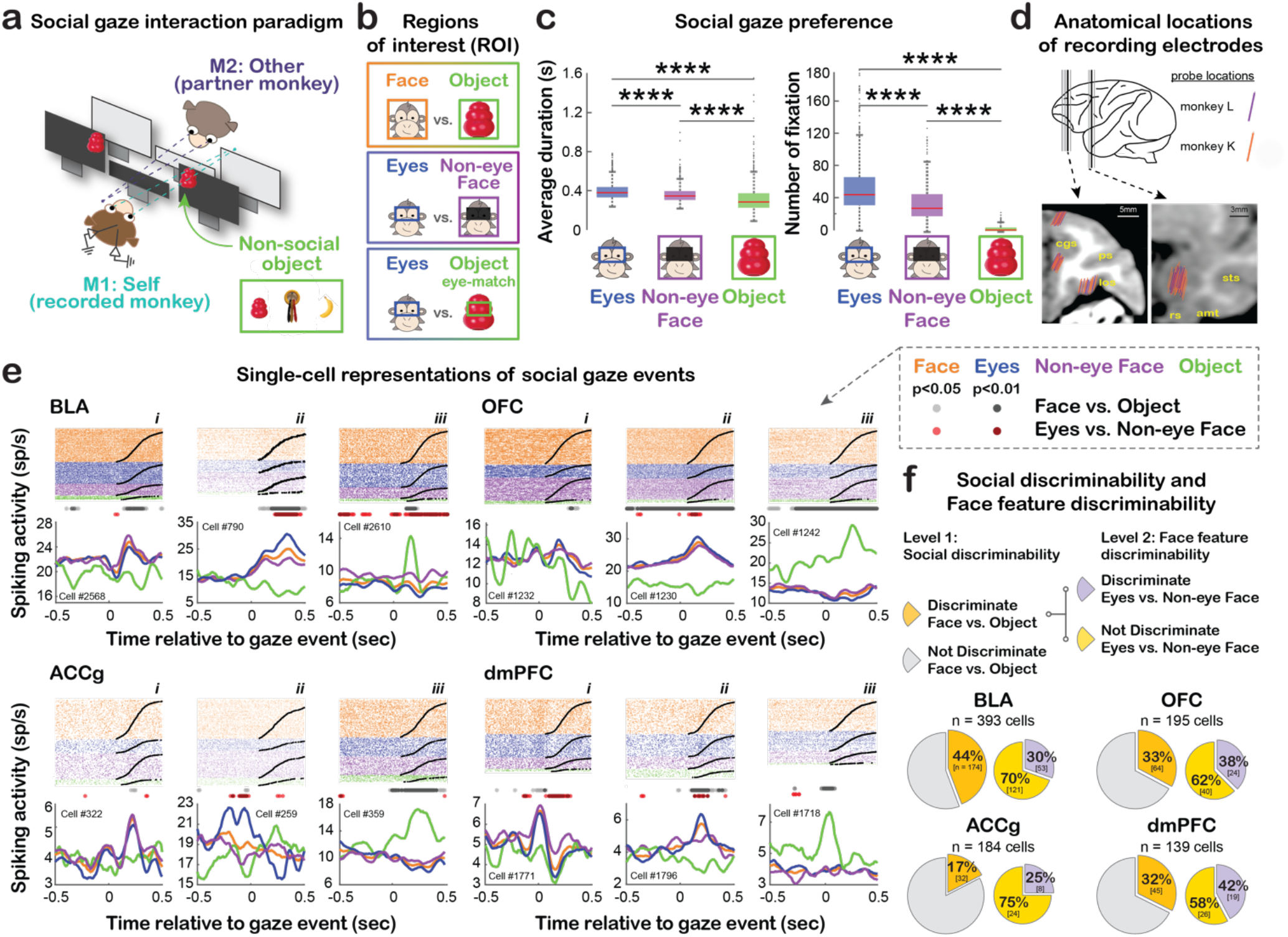
Single-cell representations of social gaze events during live social gaze interactions. **a**, Experimental paradigm for studying naturalistic, face-to-face, social gaze interaction between pairs of monkeys. The inset shows three different types of non-social objects used. **b**, Illustrations of the four gaze ROIs and contrasts. **c**, Social gaze preference, indicated as the average duration per fixation and the total number of fixations to *Eyes*, *Non-eye Face* and *Object*. ****, p < 0.0001, Wilcoxon sign rank, two-sided, FDR-corrected. **d**, Anatomical locations of electrode positions from monkeys L and K shown on representative coronal MRI slices of monkey K (Fig. S2 shows individual cell locations for both monkeys) (cgs, cingulate sulcus; ps, principal sulcus; los, lateral orbitofrontal sulcus; sts, superior temporal sulcus; amt, anterior middle temporal sulcus; rs, rhinal sulcus). **e**, Single-cell examples of spiking activity around social gaze events. For each cell, spike rasters are shown at the top, and the peristimulus time histogram (PSTH) shows the average firing rate aligned to gaze event onset (orange: *Face*; blue: *Eyes*; purple: *Non-eye Face*; green: *Object*). Light gray (p < 0.05, hierarchical ANOVA) and dark gray (p < 0.01) circles indicate time bins with significantly different activity for *Face* vs. *Object*. Light red (p < 0.05) and dark red (p < 0.01) circles indicate time bins with significantly different activity for *Eyes* vs. *Non-eye Face*. **f,** Hierarchical classification of cells with ‘social discriminability’ (*Face* vs. *Object*) and ‘face feature discriminability’ (*Eyes* vs. *Non-eye Face*). Larger pies show the proportions of cells in each area with (dark yellow) and without (gray) ‘social discriminability’. BLA showed the highest proportion of cells with ‘social discriminability’ (vs. OFC: ***χ***^**2**^ = 7.10, p < 0.01; vs. ACCg: ***χ***^**2**^ = 39.46, p < 0.0001; vs. dmPFC: ***χ***^**2**^ = 6.00, p = 0.02, Chi-square test, FDR-corrected). By contrast, ACCg showed the lowest proportion of cells with ‘social discriminability’ (vs. OFC: ***χ***^**2**^ = 11.92, p < 0.001; vs. dmPFC: ***χ***^**2**^ = 9.79, p < 0.005, FDR-corrected). Smaller pies show the proportions of cells with ‘social discriminability’ that further showed (purple) and did not show (light yellow) ‘face feature discriminability’. BLA, OFC, and ACCg had higher number of cells with only ‘social discriminability’ compared to cells with further ‘face feature discriminability’ (all ***χ***^**2**^ > 8.00, p < 0.01, FDR-corrected), while these proportions were comparable in dmPFC (***χ***^**2**^ = 2.18, p = 0.16, FDR-corrected).

During these gaze interactions, we examined spiking activity from 537 BLA, 241 OFC, 236 ACCg, and 187 dmPFC cells (Fig. 1d, Fig. S2). All four brain regions contained a considerable number of cells that categorically fired more for looking at *Face* compared to *Object* while showing indifferent activity between *Eyes* and *Non-eye Face* (Fig. 1e, subpanels *i*). Another population of cells, again in all four areas, further differentiated *Eyes* from *Non-eye Face* by either having higher or lower activity for one or the other ROI (Fig. 1e, *ii*). Finally, a third group of cells showed higher activity for *Object* than social ROIs (Fig. 1e, *iii*).

In all four neural populations, we observed considerable proportions of cells exhibiting ‘social discriminability’ (distinct activity for *Face* versus [vs.] *Object*) – 44% of BLA, 33% of OFC, 17% of ACCg, and 32% of dmPFC cells (hierarchical ANOVA; Methods) (Fig. 1f) (see Fig. S3 for the analyses controlling for the location of previous gaze fixation and central fixation). Specifically, BLA had a higher proportion of such cells than the three prefrontal areas (all *χ*^2^ > 6.00, p < 0.02, Chi-square test, FDR-corrected). Among cells with ‘social discriminability’, we observed comparable proportions of cells that displayed ‘face feature discriminability’ (distinct activity for *Eyes* vs. *Non-eye Face*) across BLA (30%), OFC (38%) ACCg (25%), and dmPFC (42%) (Fig. 1f; all *χ*^2^ < 2.44, p > 0.40, FDR-corrected). Overall, all four brain areas contained robust single-cell correlates of social gaze events with BLA showing the highest proportion of cells with ‘social discriminability’.

### Temporal diversity of social gaze encoding during live social gaze interaction

In each brain region, the times at which different cells began to differentiate gaze events varied greatly – some started to show distinct anticipatory activity for different ROIs leading up to the onset of gaze events, while others displayed distinct evaluative activity once the gaze events had occurred. All four brain areas exhibited such temporally heterogeneous activity for looking at *Face* vs. *Object* (Fig. 2a), *Eyes* vs. *Non-eye Face* (Fig. 2b), and *Eyes* vs. *Object* (Fig. S4a), suggesting temporally diverse contributions of different cells in discriminating gaze events. Specifically, higher proportions of BLA and dmPFC cells started to discriminate *Face* from *Object* in the pre-gaze compared to post-gaze epoch (Fig. 2a; both *χ*^2^ > 9.29, p < 0.005, Chi-square test, FDR-corrected), suggesting a bias towards an anticipatory processing compared to an evaluative processing for looking at *Face* vs. *Object* specifically in these two areas.

**Figure 2.**
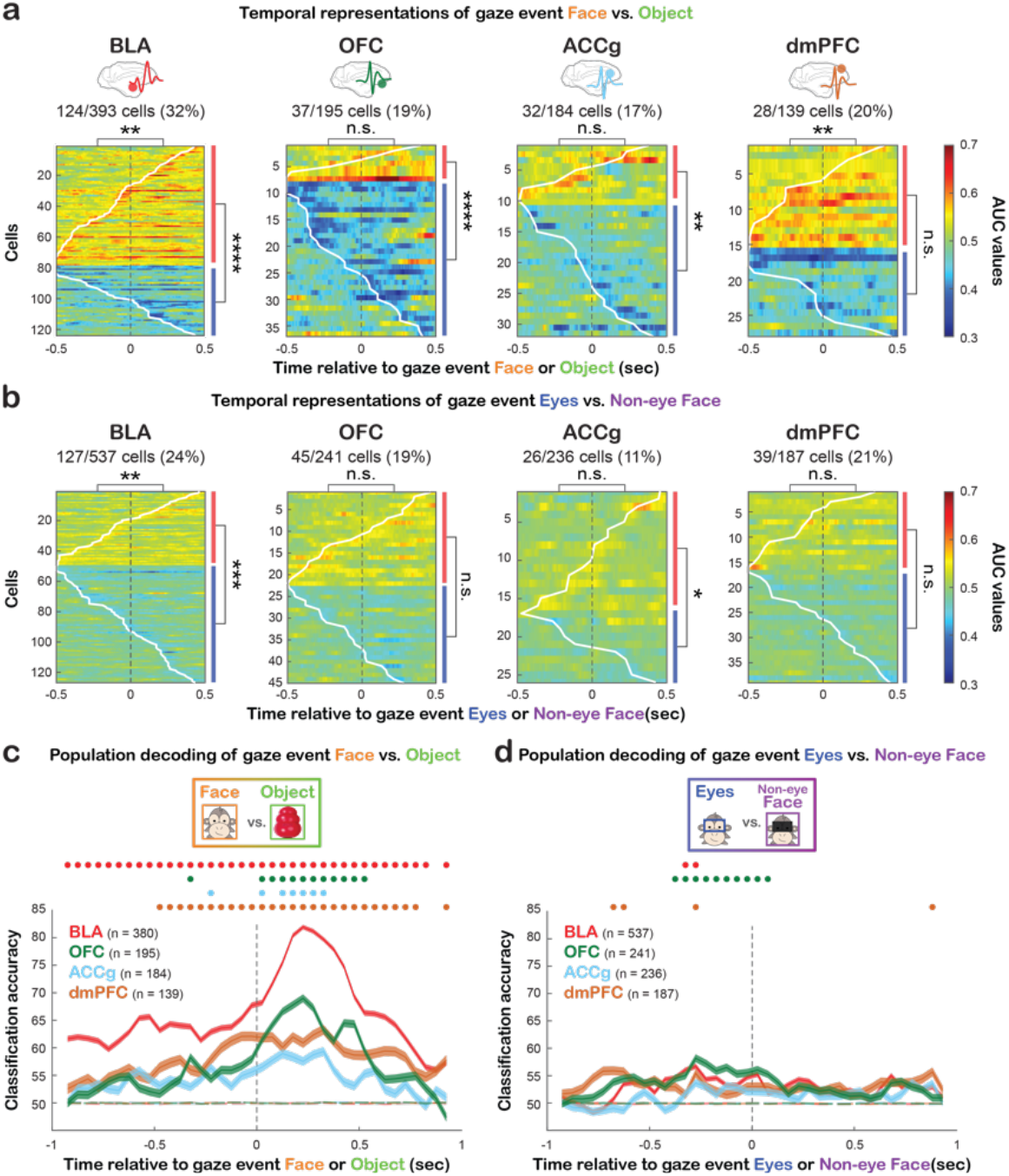
Temporal representations of social gaze events during live social gaze interaction. **a**, Temporal representations for discriminating *Face* vs. *Object* with matching ROI size. Heatmaps show the area under the curve (AUC) values from receiver operating characteristic analysis (ROC) of individual cells in BLA, OFC, ACCg, and dmPFC for significantly discriminating *Face* from *Object*. The data are aligned to the time of gaze event onset with each row representing a cell sorted based on the first bin with significant AUC (white contour). Warm colors indicate greater activity for looking at *Face* (AUC > 0.5), whereas cold colors indicate greater activity for *Object* (AUC < 0.5) (Methods). To the right of each heatmap, the red and blue bars represent the proportion of cells with greater activity for *Face* and *Object*, respectively (Methods). The asterisks on the top of each heatmap indicate the comparisons of the proportions of cells that began discriminating *Face* vs. *Object* during the pre-gaze versus post-gaze epoch. **, p < 0.01; ****, p < 0.0001; n.s, not significant, Chi-square test, FDR-corrected. **b**, Temporal profiles of spiking activity for *Eyes* vs. *Non-eye Face* (same format as **a**). **c,** Population decoding accuracy for *Face* vs. *Object* in BLA (red), OFC (green), ACCg (blue), and dmPFC (brown). Real data are shown in solid lines and empirically derived null data are in dotted lines. Circles at the top indicate the time bins with decoding accuracy significantly higher than the null in corresponding colors (p < 0.001, permutation test). **d**, Population decoding accuracy for *Eyes* vs. *Non-eye Face* (same format as **c**).

Neuronal activity in these brain regions could be used to selectively discriminate gaze events. By training a max correlation coefficient (MCC) pattern classifier (Methods), we observed significantly higher decoding accuracy for discriminating *Face* from *Object* than the empirical null in all four regions (Fig. 2c). Especially, BLA and OFC reached significant decoding accuracy greater than 80% and 65%, respectively, soon after the onset of gaze events. Furthermore, BLA and dmPFC showed significant decoding accuracy even before gaze event onsets, which is consistent with the bias towards an anticipatory processing based on AUC measures (Fig. 2a). Notably, while we observed significant decoding accuracy in BLA, OFC, and dmPFC for *Eyes vs. Object* (Fig. S4b), decoding accuracy was overall near chance in all regions for *Eyes vs. Non-eye Face* (Fig. 2d), endorsing a predominant ‘social discriminability’ in these neurons over ‘face feature discriminability’ during live social gaze interaction. Comparable decoding results were observed when the analyses were performed using the same number of cells across regions (Fig. S5). Overall, while cells in all four regions displayed temporal heterogeneity around gaze events, BLA and dmPFC showed a temporal bias toward an anticipatory processing. Moreover, the decoding performance was noticeably higher in BLA, OFC, and dmPFC relative to ACCg, suggesting that these three populations exhibited more robust ‘social discriminability’.

### Social gaze monitoring from the perspective of self, other, or both during live social gaze interaction

To understand how individual cells modulated spiking activity across the visual environment surrounding a conspecific or an object, we constructed a spike density map using M1’s gaze positions relative to the center of M2’s *Eyes*, *Face*, or *Object*. Some cells increased activity selectively when fixating within *Eyes* or *Face*, but not *Object* (Fig. S6, first column), while others did so in a more spatially distributed manner (Fig. S6, second column). Another group of cells fired more selectively for fixations closer to *Object* (Fig. S6, third column). At the population level, all four regions exhibited greater mean spiking activity for looking closer to *Face* and *Eyes*, compared to *Object* (Fig. 3a–b; ROI main effect, F(2, 3293) = 885.47, p < 0.0001, two-way ANOVA), such that all four regions exhibited greater mean activity for *Face* and *Eyes*, than *Object* (Fig. 3b, all p < 0.0001, Tukey test), resulting in spike density maps focally centered on the social stimuli (Fig. 3a).

**Figure 3.**
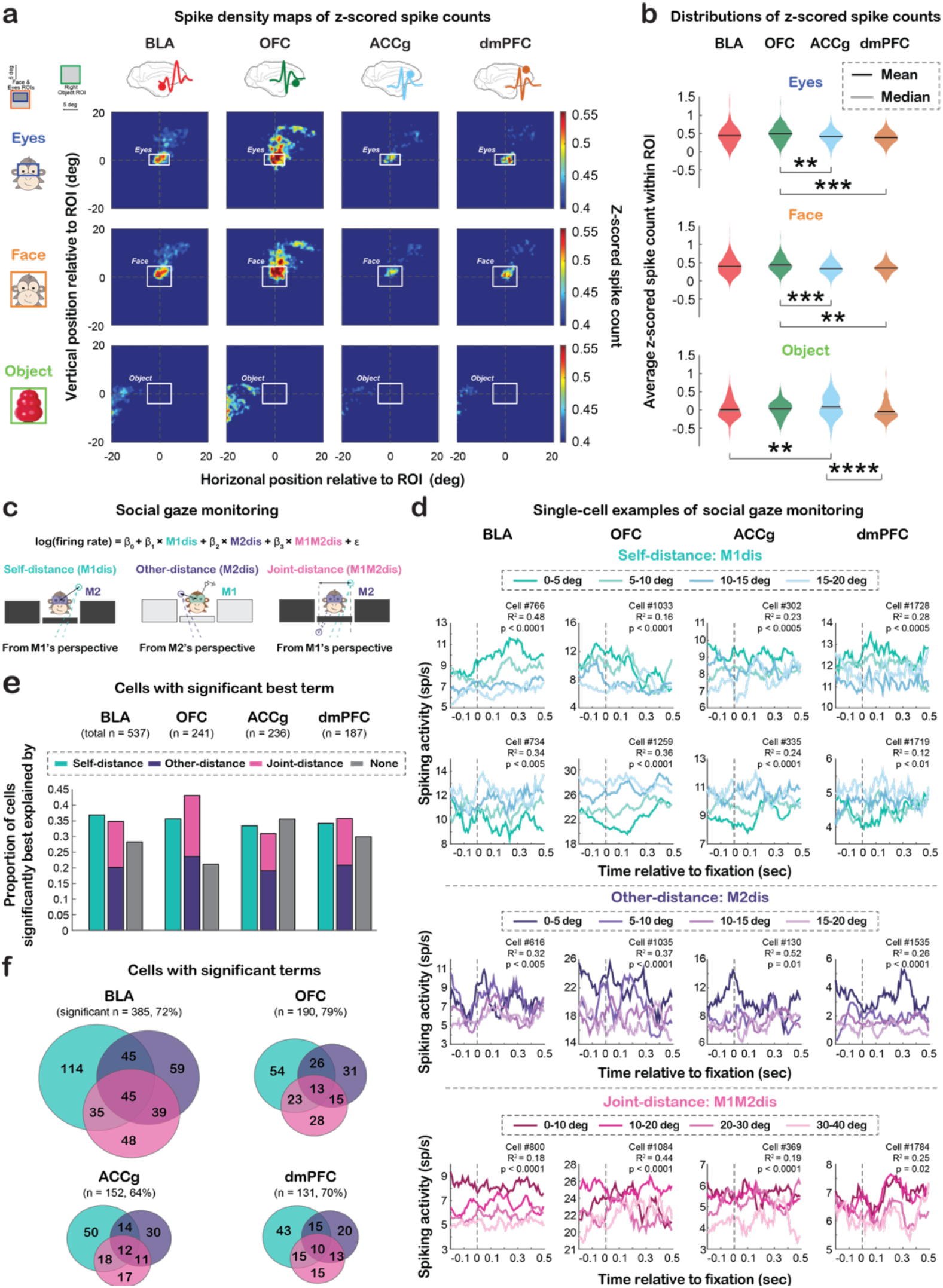
Social gaze monitoring from the perspective of self, other, or both (joint). **a**, Spike density maps (z-scored spike counts) aligned to the center of partner’s *Eyes* (top), *Face* (middle) and *Object* (bottom; matching ROI size as *Face*) in BLA, OFC, ACCg and dmPFC populations. Top left inset, the layout of the setup for reference. **b**, Distributions of the z-scored spike counts. All four regions exhibited greater activity for *Eyes* and *Face*, compared to *Object* (see text). Compared to dmPFC and ACCg, OFC population showed greater activity for fixating *Eyes* and *Face* (all p < 0.005, Tukey test), while ACCg showed greater activity for fixating within *Object* compared to BLA and dmPFC (both p < 0.005, Tukey test). **, p < 0.01; ***, p < 0.001; ****, p < 0.0001, two-way ANOVA, Tukey test. **c**, Modeling of social gaze monitoring with illustrations of the three distance variables involved: Self-distance (*M1dis*, mint), Other-distance (*M2dis*, purple), and Joint-distance (*M1M2dis*, pink) (Eq. 1–2; Methods). **d**, Single-cell PSTH examples from each area whose activity was significantly best explained by Self-distance (top; showing two modulation types), Other-distance (middle), or Joint-distance (bottom) (see Fig. S7 for the distributions of modulation types). **e**, Proportions of cells in each area whose activity was significantly best explained by Self-distance, Other-distance, Joint-distance, or none (gray). **f**, Venn diagrams of cells in each region whose activity could be significantly explained by one or more of the three distance variables, independent of the best-fitting term.

These results raised an intriguing possibility that neurons in these regions might track one’s gaze positions referenced to another agent, such that cells may fire more when looking closer toward and less when looking farther away from a conspecific. We thus tested if activity parametrically tracked the gaze positions of self (M1) or other (M2) in a continuous manner by examining three social gaze-related distance variables – Self-distance (the distance between M1’s gaze positions and the center of M2’s eyes), Other-distance (the distance between M2’s gaze positions and the center of M1’s eyes), and Joint-distance (the distance between the gaze positions of the two monkeys) (Methods; Eq. 1–2) (Fig. 3c).

In all four regions, activity of many cells was significantly explained by Self-distance, Other-distance, or Joint-distance, where activity either increased or decreased as distance variables increased (Fig. 3d, Fig. S7a). Activity of 37% of BLA, 36% of OFC, 33% of ACCg, and 34% of dmPFC cells was significantly best explained by Self-distance (Fig. 3e). Notably, activity of many cells in each region was best explained by either Other-distance or Joint-distance, the two variables considering the gaze positions of the partner monkey – 35% of BLA, 43% of OFC, 31% of ACCg, and 36% of dmPFC (Fig. 3e). When considering all significant terms for each cell, regardless of the best-fitting term, we observed numerous cells that concurrently tracked more than one distance variable (Fig. 3f). Similar modeling results were observed by using a subset of cells with relatively higher variance explained (Fig. S7b–c; Methods). Specifically, considering all cells, there was a regional difference in the proportions of cells tracking at least one distance variable (*χ*^2^ = 12.38, p = 0.022, FDR-corrected), driven by more such cells in OFC than ACCg (*χ*^2^ = 11.54, p < 0.005, FDR-corrected).

Modeling of social gaze monitoring using these distance variables provided remarkably good fits as the true mean adjusted *R*^*2*^ across all cells in each area was always greater than the null distributions (Fig. S7d; all p = 0, permutation test; Methods). The modeling results were also relatable to the AUC results (Fig. 2) such that neural discriminability of other’s *Eyes* from *Object* was systematically correlated with the tracking of Self-distance (Fig. S7e; rho = −0.31 and −0.44 for BLA and OFC, respectively, both p < 0.0005, rho = −0.43, p < 0.001 for ACCg, and rho = −0.36, p = 0.02 for dmPFC; Spearman correlation). Overall, the neural tracking of Self-distance, Other-distance, or Joint-distance supports a single-cell mechanism of social gaze monitoring in the four brain regions.

### Mutual eye contact selectivity during live social gaze interaction

An important component of social gaze interaction is mutual eye contact, defined as when both individuals look at each other’s eyes simultaneously (Fig. 4a). To understand the neuronal correlates of these interactive gaze behaviors, we examined *Interactive Mutual Eyes* events (Fig. 4b–c), as a function of two separate agent-specific contexts: *Self-follow Mutual Eyes* (i.e., M2 looked at M1’s *Eyes*, followed by M1 looking at M2’s *Eyes*) or *Other-follow Mutual Eyes* (M1 looked at M2’s *Eyes*, followed by M2 looking at M1’s *Eyes*). Neural activity of *Interactive Mutual Eyes* events was contrasted to the corresponding *Solo Eyes* events, defined as when only one monkey in the pair looked at the other’s eyes without any reciprocating gaze from the other (Methods). Each cell’s activity during *Self-follow Mutual Eyes* was compared to *Self Solo Eyes* (Fig. 4c, top, Fig. S8a) and activity during *Other-follow Mutual Eyes* was compared to *Other Solo Eyes* (Fig. 4c, bottom, Fig. S8b).

**Figure 4.**
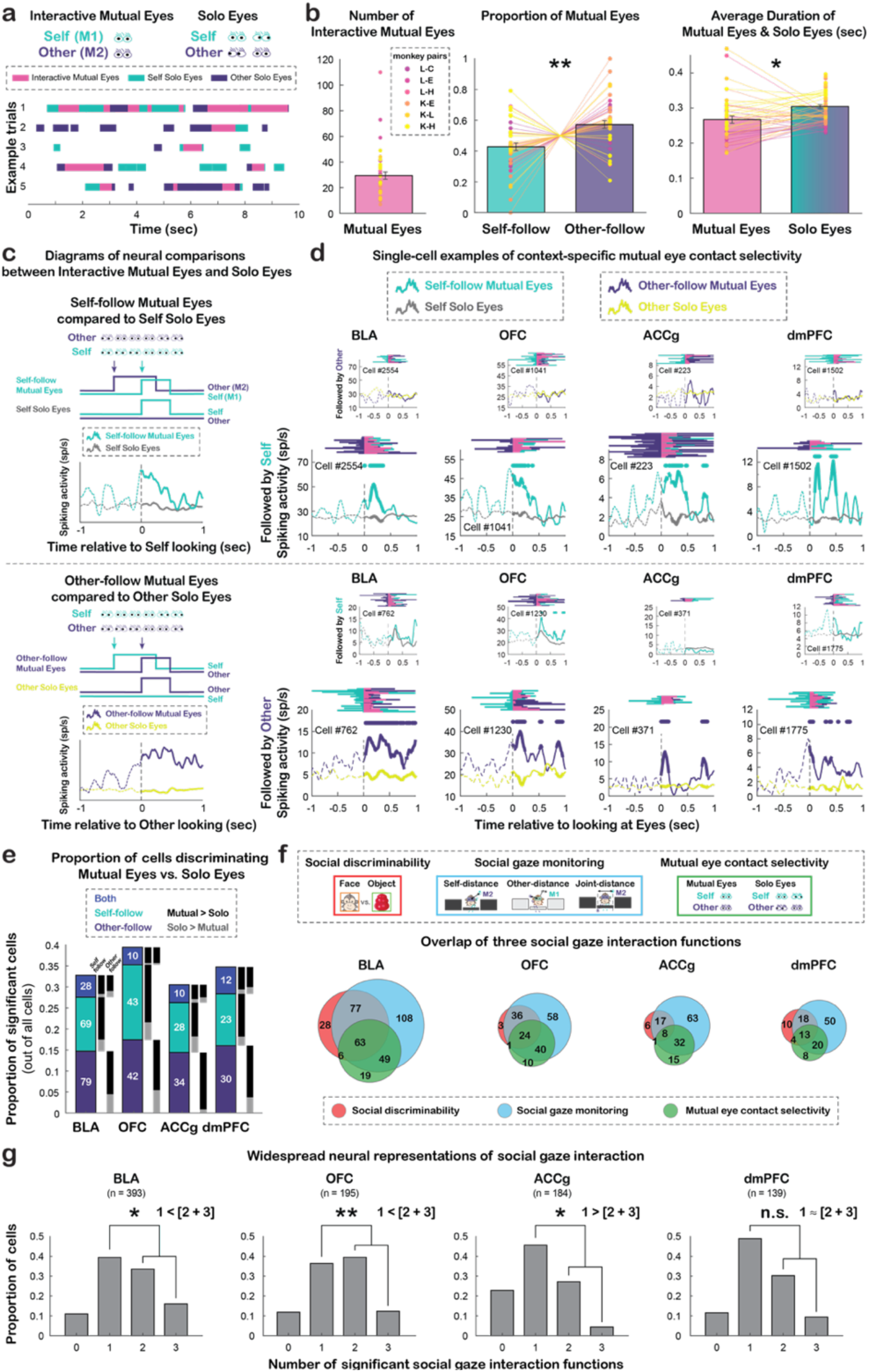
Mutual eye contact selectivity during live social gaze interactions. **a**, Examples of gaze interaction bouts between pairs of monkeys, including *Interactive Mutual Eyes* events and *Solo Eyes* events. **b**, Total number of *Interactive Mutual Eyes* events per day across all pairs of monkeys (left), proportions of *Self-follow* and *Other-follow Mutual Eyes* (middle), and average durations of *Mutual Eyes* and *Solo Eyes* events collapsed across monkeys (right). *, p < 0.05; **, p < 0.01, Wilcoxon signed rank, two-sided. **c**, Diagrams illustrating the neural comparisons between *Self-follow Interactive Mutual Eyes* (mint) and *Self Solo Eyes* (gray), aligned to the time of M1 (self) looking at M2’s (other) eyes, and between *Other-follow Interactive Mutual Eyes* (purple) and *Other Solo Eyes* (mustard), aligned to the time of M2 looking at M1’s eyes. **d**, Single-cell PSTH examples showing context-specific activity of mutual eye contact selectivity. Top, example cells from each area with selectivity for *Self-follow Mutual Eyes* (large panels), but not for *Other-follow Mutual Eyes* (small panels). Bottom, example cells from each area with selectivity for *Other-follow Mutual Eyes* (large panels), but not for *Self-follow Mutual Eyes* (small panels). For each panel, behavioral rasters are shown at the top (mint: M1 looking at M2’s eyes; purple: M2 looking at M1’s eyes; pink: mutual eye gaze). Each PSTH shows the average firing rate aligned to the time of *Interactive Mutual Eyes* events and corresponding *Solo Eyes* events. Traces in the analyzed epoch (first 500 msec following gaze event onsets) are shown as solid lines. Circles above the traces indicate time bins with significantly different activity between *Interactive Mutual Eyes* and *Solo Eyes* events (p < 0.05, Wilcoxon rank sum test, two-sided). **e**, Proportions of significant cells out of all cells from each area that selectively differentiated *Self-follow Mutual Eyes* from *Self Solo Eyes* (mint), selectively differentiated *Other-follow Mutual Eyes* from *Other Solo Eyes* (purple), or differentiated both types of comparisons (blue). The black bars indicate the proportions of cells with greater activity for *Interactive Mutual Eyes* than *Solo Eyes*, whereas the gray bars indicate the opposite. **f–g**, Proportions of cells showing correlates of one or more of the three social gaze interaction functions, namely, social discriminability (red), social gaze monitoring (blue), and mutual eye contact selectivity (green). **f,**Venn diagrams summarize the single-cell level overlaps among the three gaze functions for each brain region. **g**, Bar graphs show the proportion of cells in each area that were involved in none, one, two, or all three of the social gaze interaction functions. *, p = 0.01; **, p < 0.01, n.s., not significant, Chi-square test, FDR-corrected.

In all four brain regions, we observed cells that differentiated *Interactive Mutual Eyes* from *Solo Eyes*. Some cells exclusively differentiated *Self-follow Mutual Eyes* from *Self Solo Eyes* (Fig. 4d, top), while others exclusively differentiated *Other-follow Mutual Eyes* from *Other Solo Eyes* (Fig. 4d, bottom), supporting selective representations of agent-specific mutual eye contact context. In all brain areas, similar proportions of cells were selective for *Self-follow Mutual Eyes* or *Other-follow Mutual Eyes* (Fig. 4e; all *χ*^2^ < 0.93, p > 0.33, Chi-square test, FDR-corrected), while a significantly smaller group of cells (Fig. S9a) signaled both types of mutual eyes events (Fig. 4e; both types vs. *Self-follow*: *χ*^2^ > 8.53, p < 0.005 for BLA, OFC, and ACCg, *χ*^2^ *=* 3.46, p = 0.06 for dmPFC; both types vs. *Other-follow*: all *χ*^2^ > 7.71, p < 0.01; FDR-corrected). Crucially, cells in all brain areas mostly showed higher activity for *Interactive Mutual Eyes* than *Solo Eyes* (all *χ*^2^ > 5.14, p < 0.03 for 12 of 16 cases, FDR-corrected). Similar results were obtained at a later time epoch (500–1000 msec following the gaze event onsets) (Fig. S9b) or only using the subset of cells with ‘social discriminability’ (Fig. S9c). Overall, many cells in these four areas displayed selectivity for agent-specific mutual eye contact context, and OFC had the highest proportion of cells with selectivity for mutual eye contact.

Overall, we observed widespread single-cell implementations of social gaze interaction within the primate social interaction networks (Freiwald, 2020) – neuronal correlates of social discriminability, social gaze monitoring, and mutual eye contact selectivity were broadly found across BLA, OFC, ACCg, and dmPFC (Fig. 4f). Remarkably, at least 77% and 31% of cells from each brain region significantly signaled at least one and two of the three social gaze interaction functions examined, respectively (Fig. 4g). Such extensive representations are likely indicative of the evolutionary pressure put on the primate brain for regulating complex social exchanges among individuals.

There were also significant interareal differences, such that a greater proportion of BLA cells displayed ‘social discriminability’ than the three prefrontal areas, with ACCg containing the least proportion of such cells (Fig. 1f). Further, BLA and dmPFC showed a temporal bias for an anticipatory processing of looking at *Face* vs. *Object* (Fig. 2a), while ACCg contained the least decodable information (Fig. 2c). Moreover, a significantly greater proportion of OFC than ACCg cells was involved in social gaze monitoring (Fig. 3f), and a particularly high proportion of OFC neurons showed selectivity for mutual eye contact (Fig. 4e). Important future research directions include understanding how multiple brain regions with shared functionalities within the social interaction networks(Freiwald, 2020) work together to enable social interactions.

Intriguingly, some neurons in these regions only signaled one of the three social gaze interaction functions examined, suggesting that there might be functional specializations, whereas other cells signaled at least two of the three functions examined (Fig. 4f–g), endorsing a potentially shared coding schema linking them in overlapping populations. Although there was no regional difference in the proportion of cells displaying the correlates of only one function (Fig. 4g; *χ*^2^ = 7.23, p = 0.09, Chi-square test, FDR-corrected), the proportions of cells displaying the correlates of at least two functions were different across regions (*χ*^2^ = 22.19, p < 0.0005, FDR-corrected). Specifically, more BLA and OFC cells displayed correlates of at least two functions (compared to only one function: BLA, *χ*^2^ = 7.83, p = 0.01; OFC, *χ*^2^ = 8.75, p < 0.01, FDR-corrected), whereas more ACCg cells showed correlates of only one function (compared to at least two functions: *χ*^2^ = 7.17, p = 0.01, FDR-corrected) with no such difference in dmPFC (*χ*^2^ = 2.10, p = 0.16, FDR-corrected). Taken together, BLA and OFC overall showed evidence for more robust involvements in social gaze interaction compared to the other two areas, based on the three social gaze interaction signatures we examined. Recent advances have shed light on whether a shared coding is applied for multiple functions. For example, many cells in the macaque amygdala were found to use a value-based coding schema for human intruder’s gaze direction (Pryluk et al., 2020) and for social rank(Munuera et al., 2018). By contrast, evidence in mice suggests that distinct neuronal ensembles are recruited between social and non-social processing or even between different types of social processing (Allsop et al., 2018; Jennings et al., 2019; Kingsbury et al., 2020). Our finding suggests that social gaze interaction recruits partially overlapping neuronal ensembles, thereby resulting in some of the overlapping functionalities among the three social gaze interaction variables.

Complex social gaze interaction is a hallmark of primates, including perception of social cues and sophisticated interpersonal communication. Neural systems involved in social perception are distributed across the temporal and visual cortical areas (Haxby et al., 2000), including the hierarchically modular face patches in the inferior temporal cortex (Freiwald and Tsao, 2010; Koyano et al., 2021; Leopold et al., 2006; McMahon et al., 2015; Tsao, 2006), a prefrontal face patch in OFC (Barat et al., 2018; Rolls et al., 2006; Tsao et al., 2008), and the human fusiform gyrus (Kanwisher et al., 1997; McCarthy et al., 1997). When might such perceptual signal begin to transition to more action-oriented signals used for social interaction? A recent work found that a gaze-following patch in the superior temporal sulcus might mediate one of such behavioral transition from perception to interaction (Marciniak et al., 2014; Ramezanpour and Thier, 2020), suggesting an action-oriented organization of social information in certain brain regions. Moreover, neurons in the amygdala signal the perception of facial expression and mutual eye gaze (Gilardeau et al., 2021; Gothard et al., 2007; Livneh et al., 2012; Mosher et al., 2014; Rutishauser et al., 2011; Wang et al., 2017), two processes that are intricately linked to interacting with others. Notably, recent neuroimaging work in macaques demonstrated specific brain activations for the interactive aspects of social behaviors widely spanning across multiple brain areas (Shepherd and Freiwald, 2018; Sliwa and Freiwald, 2017). Moreover, accumulating evidence supports critical roles of the prefrontal-amygdala circuits in guiding social behaviors across humans, non-human primates, and rodents (Gangopadhyay et al., 2021). In the present study, we reported how single neurons in the prefrontal-amygdala circuits within the primate social interaction network (Freiwald, 2020) encode social gaze interaction, providing insights into single-cell foundations of primate social interaction and supporting that distributed and globally integrated networks guide social gaze interaction.

## Acknowledgements

This work was supported by the National Institute of Mental Health (R01MH110750, R01MH120081). We are extremely grateful to Katalin Gothard, Daeyeol Lee, Rony Paz, and Bijan Pesaran for providing helpful feedback about this work at multiple occasions. We also thank James McPartland for the discussions on the clinical relevance of our findings in autism. Finally, we thank Olivia Meisner for helpful comments on the manuscript.

## Author Contributions

S.W.C.C. and O.D.M. designed the study. O.D.M. and S.F. collected the data. O.D.M., S.F., N.A.F., C.J.C., M.B.Z., P.T.P., and S.W.C.C. analyzed the data. S.W.C.C., S.F., O.D.M., N.A.F., and A.R.N. wrote the paper.

## Data & Code Availability

Behavioral and neural data as well as the main analysis codes relevant for this paper will be available upon the publication of this work.

## Methods

### Animals

Two adult male rhesus macaques (*Macaca mulatta*) were involved as recorded monkeys (M1; monkeys L and K; aged 8 and 7 years, weighing 15.7 kg and 10 kg, respectively). Two other adult male and one adult female monkeys (monkeys C, H, and E, all aged 7 years, weighing 10.1kg, 11.1kg, and 10.7kg, respectively) served as partner monkeys (M2) during live social gaze interactions, resulting in six distinct macaque pairs (monkeys L-C, L-E, L-H, K-E, K-L, K-H). The recorded and partner monkeys were unrelated and were housed in the same colony room with other macaques. The current data collection was focused on investigating single-cell activity during spontaneous, face-to-face social gaze interaction and did not have the necessary number of pairs to examine the modulatory effects of social relationship, such as social rank. Within the same-sex pairs, monkey L was dominant over monkey C but subordinate to monkey H, whereas monkey K was dominant over monkey L but subordinate to monkey H. Our previously published work (Dal Monte et al., 2016) using the identical paradigm provides a comprehensive examination of social relationship effects on social gaze interaction from unique 8 dominance-related, 20 familiarity-related, and 20 sex-related perspectives. All animals were kept on a 12-hr light/dark cycle with unrestricted access to food, but controlled access to fluid during testing. All procedures were approved by the Yale Institutional Animal Care and Use Committee and in compliance with the National Institutes of Health Guide for the Care and Use of Laboratory Animals. No animals were excluded from our analyses.

### Surgery and anatomical localization

All animals received a surgically implanted headpost (Grey Matter Research) for restraining their head movement. A second surgery was performed on the two recorded animals to implant a recording chamber (Crist and Rogue Research Inc.) to permit neural recordings from BLA, OFC (Brodmann areas 11 and 13m), ACCg (24a, 24b and 32), and dmPFC (8Bm and F6) (Paxinos et al., 1999) (see Fig. 1d for the summary of electrode locations on representative MR slices; see Fig. S2 for the locations of individual cells on the Paxinos slices). Placement of the chambers was guided by both structural magnetic resonance imaging (MRI, 3T Siemens) scans and stereotaxic coordinates.

### Experimental setup

On each day, M1 and M2 sat in primate chairs (Precision Engineering, Inc.) facing each other, 100 cm apart with the top of each monkey’s head 75 cm from the floor (Fig 1a, Fig. S1a–b). Each monkey faced three monitors with the middle monitor 36 cm away from the eyes. Two infrared eye-tracking cameras (EyeLink 1000, SR Research) continuously and simultaneously recorded the horizontal and vertical eye positions of both monkeys.

Each monkey first underwent a standard eye position calibration procedure. The middle monitor displayed five stimuli in different locations, controlled by Psychtoolbox (Brainard, 1997) and EyeLink toolbox (Cornelissen et al., 2002) in MATLAB. Each monkey was required to fixate on these stimuli sequentially to calibrate and register eye positions. During this procedure, neither monkey had visual access to the other. Critically, because each animal’s face was on a different depth plane from the monitors, we carried out an additional calibration procedure to precisely map out each monkey’s facial regions. To do so, we designed a customized face calibration board (23 cm L×18 cm H×1.5 cm W) embedded with LED lights that were aligned to each monkey’s eyes, mouth, and the four corners of the face (Fig. S1c–e). This custom board was first positioned in front of a monkey’s face, secured on the primate chair, and then the middle monitors were lowered down remotely using a controlled hydraulic system. The monkey undergoing this calibration was required to fixate on these LED lights in sequence to register eye positions, after which the middle monitors were raised up to block the view of the two monkeys. The same procedure was immediately repeated for the second monkey. The middle monitors then remained raised up until the beginning of recording sessions.

Each recording day consisted of a total of 10 social gaze interaction sessions. At the beginning of each session, the middle monitors were lowered down remotely so that the two monkeys could fully see each other (Fig 1a, Fig. S1a–b). During each session, monkeys could freely make spontaneous eye movements and interact with each other using gaze for five minutes. At the end of each 5-min session, the middle monitors were raised up remotely, and monkeys had no visual access of one another during a 3-min break (inter-session breaks). On M1’s side, two identical objects (chosen from monkey toy cones, toy keys, or bananas, on different days) were attached to the monitors at 20.7° eccentricity throughout the sessions to serve as non-social objects (Fig 1a).

### Gaze regions of interest

We identified four gaze regions of interest (ROIs): *Face*, *Eyes*, *Non-eye Face* (i.e., face area excluding the eyes), and *Object* (Fig 1b). From each day’s calibration, the *Face* ROI was defined by the four corners of a monkey’s face, and the *Eyes* ROI was defined by adding a padding of 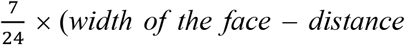 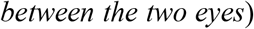 to the center of each eye. The *Object* ROI had the same surface area as the *Face* ROI, unless when it was directly compared to the *Eyes* ROI where we matched its surface area to that of the *Eyes* ROI (Fig. 1b). The EyeMMV toolbox (Krassanakis et al., 2014) in MATLAB was used to detect fixations based on spatial and duration parameters. A gaze event was defined as a fixation within a specific ROI with a minimum duration of 70 msec. For the three non-overlapping ROIs, *Eyes*, *Non-eye Face*, and *Object*, we calculated the total number of M1’s fixations and the average duration of each fixation for each day. One-way ANOVA was used to compare each variable across the three ROIs (Fig 1c).

### Single-unit activity

Single-unit activity was recorded from 16-channel axial array electrodes (U-or V-Probes, Plexon Inc.) using a 64-channel system (Plexon Inc.) (Fig. 1d). A guide tube was used to penetrate intact dura and to guide electrodes, which were remotely lowered by using a motorized multi-electrode microdrive system (NaN Instruments) at the speed of 0.02 mm/sec. After electrodes reached targeted sites, we waited 30 min for the tissue to settle and to ensure signal stability before starting the neural recording.

Broadband analog signals were amplified, band-pass filtered (250 Hz–8 kHz), and digitized (40 kHz) using a Plexon OmniPlex system. Spiking data were saved for waveform verifications offline and automatically sorted using the MountainSort algorithm (Chung et al., 2017). This resulted in a total of 537 BLA, 241 OFC, 236 ACCg, and 187 dmPFC units from two recorded monkeys (monkey L and monkey K: 225 and 312 BLA cells, 102 and 139 OFC cells, 109 and 127 ACCg cells, and 92 and 95 dmPFC cells, respectively). Peri-stimulus time histogram (PSTH) of each cell was constructed by binning spike train with 10-msec time bins and smoothing the average firing rate with 100-msec sliding windows in 10-msec steps (Fig. 1e).

### Hierarchical classification of spiking activity

For each neuron, we calculated its average firing rate during the pre-gaze epoch (500 msec leading up to the time of gaze event onset) and post-gaze epoch (500 msec following the time of gaze event onset) and applied hierarchical ANOVA for classification (Fig. 1f). We first calculated the percentage of cells with ‘social discriminability’ whose activity significantly discriminated *Face* from *Object* in either time epoch (p < 0.05).

Among these significant cells, we further calculated the percentage of cells with ‘face feature discriminability’ whose activity discriminated *Eyes* from *Non-eye Face* in either time epoch (p < 0.05). For this and further analyses where *Object* was involved in any certain pairwise comparisons, we used a subset of neurons from days when non-social objects were included in the experiment, resulting in 393 BLA, 195 OFC, 184 ACCg, and 139 dmPFC neurons from two recorded monkeys (monkey L and monkey K: 225 and 168 BLA cells, 102 and 93 OFC cells, 109 and 75 ACCg cells, and 92 and 47 dmPFC cells, respectively).

### Previous gaze fixation location control

This control analysis assessed the effects of previous gaze fixation location on the neural activity around a current *Face* event. We defined a 9-cell (3 × 3) space grid aligned to M2’s face, where the center grid cell was defined as the *Face* ROI with the remaining cells distributed around it (Fig. S3a). Each grid cell was numbered, where the *Face* ROI was associated with grid index 5, as shown in Fig. S3a. We first examined the 500 msec period preceding each current looking event to *Face*, identified the fixation right before the *Face* event, and assigned this fixation to one of the 9 spatial grid cells according to its average position. If the location of this preceding fixation fell outside the grid, or within the *Face* ROI, the corresponding current *Face* event was excluded from this analysis. We then averaged the spike counts of each neuron over the pre-gaze epoch and post-gaze epoch separately. For each time epoch and neuron, we then ran one-way ANOVA with grid index as the factor. For neurons with a significant main effect of grid index (p < 0.05), we computed, for each grid index, the proportion of significant Tukey post-hoc comparisons between that grid index and all others (p < 0.05). For neurons with no main effect of grid index, we set these proportions to 0. We then averaged these proportions across neurons to produce a heatmap (Fig. S3b). Lastly, for each brain region and time epoch, we plotted a Venn diagram to examine the overlap between neurons that significantly differentiated looking at *Face* vs. *Object* based on the hierarchical ANOVA (see above) and neurons that significantly differentiated space grids (Fig. S3c–d). A small overlap would mean that activity of cells with ‘social discriminability’ was marginally modulated by the location of previous fixation.

### Central fixation control

This control analysis compared neural activity between when M1 fixated on M2’s *Face* and when M1 fixated on a white central *Fixation square* (Fig. S3e) shown on the middle screen (when the middle screen was raised up during inter-session breaks). We used a subset of neurons from days when we presented the *Fixation square* stimuli (393 BLA, 178 OFC, 172 ACCg, and 132 dmPFC cells). For each cell, we computed average spike counts over the 500-msec period after the presentation of the central *Fixation square* on trials where M1 successfully fixated on the square for 300 msec. We then performed a Wilcoxon rank sum test (p < 0.05, two-sided) to compare this spike distribution to average spike counts over the 500-msec period following looking at M2’s *Face* to examine the percentage of cells per brain region that showed distinct activity for *Face* vs. *Fixation square*, both in the central gaze location of M1’s view (Fig. S3f). Lastly, for each brain region, we plotted a Venn diagram to evaluate the overlap between neurons that significantly differentiated *Face* vs. *Object* based on hierarchical ANOVA (see above) and neurons that significantly differentiated *Face* vs. *Fixation square* (Fig. S3g). A large overlap would mean that activity of cells with ‘social discriminability’ was unlikely driven by the different visual angle between looking at *Face* and *Object*, as many of these cells also showed distinct activity for *Face* and *Fixation square*, both requiring a central gaze fixation.

### Receiver operating characteristic analysis

For each brain area, we compared each neuron’s firing rate distribution for pairs of ROIs, including *Face* vs. *Object* (with a matching ROI size; Fig. 2a), *Eyes* vs. *Object* (with a matching ROI size; Fig. S4a), and *Eyes* vs. *Non-eye Face* (Fig. 2b) to perform the receiver operating characteristic (ROC) analysis. For each pairwise comparison, we binned spiking activity in consecutive 10-msec time bins, ranging from 500 msec before to 500 msec after M1’s corresponding gaze event onset. For each neuron, we then determined if a time window had a significant area under the curve (AUC) value by shuffling its firing rates and ROI labels 100 times (p < 0.01, permutation test). Neurons with significant AUC values for at least 5 consecutive bins were included in further analyses and were sorted based on the first bin with a significant AUC value.

For each pair of ROIs and brain region, we compared the proportions of cells that began discriminating one ROI from the other ROI during the pre-gaze epoch (“pre”) vs. during post-gaze epoch (“post”). A cell was assigned to the “pre” category if the first time point of at least five consecutive bins from the AUC sequence fell within the pre-gaze epoch, whereas a cell was assigned to the “post” category otherwise. For each ROI pair and brain area, we performed a Chi-square test to compare the relative proportions of “pre” vs. “post” cells.

Finally, we compared the proportions of cells that fired more for the first ROI in a pair than the second ROI (AUC > 0.5, “greater”; red bar to the right of each heatmap in Fig. 2a–b and Fig. S4a) to the proportions of cells that fired more for the second ROI than the first ROI (AUC < 0.5, “less”; blue bar). A cell was assigned to the “greater” category if the first time point of at least five consecutive bins from AUC sequence had greater activity for the first ROI, whereas a cell was assigned to the “less” category if that first time point had greater activity for the second ROI.

### Decoding analysis

For each brain region, we trained a max-correlation-coefficient (MCC) pattern classifier (Meyers, 2013; Meyers et al., 2018; Munuera et al., 2018) on neurons’ (pseudo-population) firing rate to discriminate between pairs of gaze events, including *Face* vs. *Object* with a matching ROI size (Fig. 2c), *Eyes* vs. *Object* with a matching ROI size (n = 911) (Fig. S4b), and *Eyes* vs. *Non-eye Face* (n = 1201) (Fig. 2d). We first z-scored spiking activity of each neuron from 500 msec before to 500 msec after the time of M1’s gaze event onset, and averaged firing rate with 150-msec sliding windows in 50-msec steps. The decoding procedure for the true model was run using the average of 1000 bootstrap iterations across 15 cross-validation splits. For each iteration, 14 of the 15 splits were used as the training data, and the remaining partition was used as the testing set. This procedure was repeated 15 times, thus each partition was used once as the testing set. We excluded neurons that had fewer than 15 repetitions, which resulted in 380 BLA, 195 OFC, 184 ACCg, and 139 dmPFC cells for *Face* vs. *Object*; 264 BLA, 89 OFC, 125 ACCg, and 110 dmPFC cells for *Eyes* vs. *Object*; and 537 BLA, 241 OFC, 236 ACCg, and 187 dmPFC cells for *Eyes* vs. *Non-eye Face*. To evaluate the significance of the decoding results, labels of gaze events were randomly shuffled, and the same decoding procedure was run using the average of 100 bootstrap iterations across 15 cross-validation splits. This procedure was repeated 1000 times to create a null distribution. A time bin was marked significant when the true decoding accuracy exceeded all the values in the null distribution (p < 0.001). Finally, to control for any effect of different numbers of cells across brain regions, we additionally ran the decoding analysis with the same number of neurons across regions by sub-sampling on each iteration for both the true and null models (across all four brain regions: 139 cells for *Face* vs. *Object*, 89 cells for *Eyes* vs. *Object*, and 187 cells for *Eyes* vs. *Non-eye Face*) and observed similar results (Fig. S5).

### Spike-density maps for social and non-social ROIs

To examine spike modulations with respect to the surrounding space of different ROIs, a spatial grid spanning 40 degrees of visual angle in both horizontal and vertical dimensions was constructed, centered on the *Eyes*, *Face*, and *Object* ROIs, separately, with 100 bins in each dimension. Each M1’s fixation was assigned to a grid-square based on the centroid of that fixation. For each neuron, the total number of spikes occurring within the 500 msec after each fixation onset was calculated and assigned to the corresponding grid-square. Total spike counts were calculated by summing across all fixations in each grid-square and were z-scored and averaged across neurons (Fig. 3a, Fig. S6). Lastly, we calculated the average z-scored spike count within each ROI and compared across brain regions and ROIs using two-way ANOVA (Fig. 3b).

### Social gaze monitoring modeling analyses

To test if and how neural activity tracks the gaze of self or other, we constructed a stepwise general linear model (GLM) for each cell. This model quantified each neuron’s firing rate in relation to three social gaze-related distance variables – 1) Self-distance (*M1dis*), the distance between recorded monkey’s gaze position and the center of partner monkey’s eyes projected to the same plane, 2) Other-distance (*M2dis*), the distance between partner monkey’s gaze position and the center of recorded monkey’s eyes projected to the same plane, and 3) Joint-distance (*M1M2dis*), the distance between recorded monkey’s gaze position and partner monkey’s gaze position projected to the same plane (Eq. 1) (Fig. 3c). We used *stepwiseglm* function in MATLAB to fit the model with a log link function. By expressing log (firing rate) as log 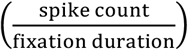, we obtained the final equation used to fit the model (Eq. 2) with the assumption that spike count follows a Poisson distribution, and set log (fixation duration) as an offset for each fixation.

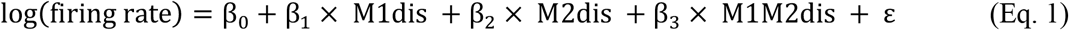

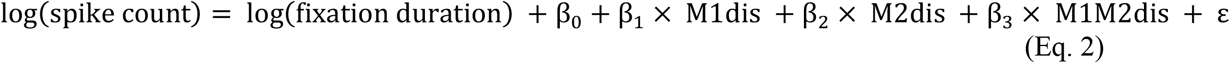

To fit the model, we first identified all of M1’s fixations in space regardless of where M2 was looking at. For each M1’s fixation, we then examined the gaze positions of M2 at 1 Hz resolution. A fixation was dropped if more than 90% of M2’s gaze samples were outside 1 degree of visual angle from the centroid of these samples during the period of that M1’s fixation. This procedure was to exclude M2’s saccades and therefore to ensure that the calculation of mean gaze position was accurate. On average, about thousand fixations were used to fit the model per neuron (mean = 1166, median = 1214). We further excluded M1’s fixation if its mean gaze position was outside 20 visual degree radius from the center of M2’s *Eyes* to reasonably ensure that M1 was able to monitor where M2 was looking at using his peripheral vision. We used 20 visual degrees based on macaque’s ability to make peripheral visual discrimination at this eccentricity in controlled visual behavioral tasks(McAlonan et al., 2008). For each of the remaining fixations, we calculated its duration, the total number of spikes during that duration, as well as the three gaze-related distance variables mentioned above – Self-distance, Other-distance, and Joint-distance. For each neuron, stepwise GLM first tested each variable and selected the one that significantly best explained the neural data. It then tested if adding a second variable would significantly improve the model performance until adding a new variable no longer improved the model. Advantages of this method are that it does not depend on the order of variable inputs and does not force a neuron to include all three variables. Therefore, the model not only informs the variables that significantly explain each neuron’s activity, but also indicates which variable does the best or worst in an order.

Single-cell PSTHs modulated by each of the three variables were constructed as in Fig. 1e (Fig. 3d). At the population level, we calculated the proportion of neurons whose spiking activity was significantly best explained by each of the three variables (Fig. 3e; p < 0.05) and examined the distribution of the coefficient of these best terms by testing if the mean coefficient was significantly different from zero for each brain region (Fig. S7a; one-sample t-test, two-sided). We further calculated the proportion of cells whose spiking activity could be significantly explained by each variable regardless of whether a variable was the best term, as long as adding the variable would significantly improve model performance (Fig. 3f; p < 0.05). Additionally, we compared the proportion of cells encoding at least one distance variable across the four brain regions. To test that our modeling results were not driven by noise in the data, we also calculated the two proportions mentioned above by using a subset of neurons with relatively high *R*^*2*^, one standard deviation greater than the mean *R*^*2*^ of all neurons across regions (mean *R*^*2*^ = 0.18; mean *R*^*2*^ + std = 0.33) (Fig. S7b–c) and observed similar results. Furthermore, to inspect the quality of model fits, we generated a null distribution of mean adjusted *R*^*2*^ by shuffling each set of Self-distance, Other-distance, and Joint-distance across fixations for 100 times for each neuron and compared the mean of this null to the observed true mean adjusted *R*^*2*^ for each brain region separately (permutation test). Lastly, to check the relationship between social gaze monitoring and social vs. non-social ROI differentiation of *Eyes* vs. *Object*, we examined Spearman correlation between the coefficient of the best term and the average AUC for *Eyes* vs. *Object* during post-gaze epoch across all neurons in each brain region (Fig. S7e).

### Mutual eye contact selectivity analyses

To examine the interactive aspects of social gaze, we focused on *Interactive Mutual Eyes* events defined as when both monkeys looked at each other’s *Eyes* simultaneously, as a function of context – that is, agent-specific sequence leading to a mutual eye contact. This resulted in two types of *Mutual Eyes* events, *Self-follow Mutual Eyes* and *Other-follow Mutual Eyes*. We compared *Mutual Eyes* events to non-interactive *Solo Eyes* events, which were defined as when only one monkey in the pair looked at the other’s eyes without any reciprocating gaze from the other monkey within at least 2-sec period around that event. To correct the discontinuity of an event when monkeys abruptly broke fixation, we smoothed the gaze vector by filling gaps between fixations less than 30 msec apart. Additionally, we excluded *Mutual Eyes* events shorter than 50 msec.

Example trials of *Interactive Mutual Eyes* and *Solo Eyes* events during social gaze interaction are shown in Figure 4a. Behaviorally, we calculated the total number of *Interactive Mutual Eyes* events per day across all pairs of monkeys (Fig. 4b, left), proportions of *Self-follow* and *Other-follow Mutual Eyes* (Fig. 4b, middle), and average durations of *Mutual Eyes* and *Solo Eyes* events collapsed across monkeys (Fig. 4b, right). To test if and how neurons discriminated *Mutual Eyes* from *Solo Eyes* events, for each neuron, we compared spiking activity associated with *Self-follow Mutual Eyes* to *Self Solo Eyes* aligned to the time of M1 looking at M2’s eyes, as well as activity associated with *Other-follow Mutual Eyes* to *Other Solo Eyes* aligned to the time of M2 looking at M1’s eyes (Fig. 4c), using two-sided Wilcoxon rank sum tests in 10-msec time bins. To reasonably ensure that the recorded monkey was able to view partner monkey in the periphery, we selected gaze events when M1’s gaze positions were within 20 visual degree radius from the center of M2’s *Eyes* ROI at least once during the time when M2 looked at M1 (noted as ‘M1 gaze criterion’ in Fig. S8). For example, for *Self-follow Mutual Eyes,* we analyzed events when M1’s gaze positions were within 20 visual degrees at any time point during the time when M2 was looking at M1’s eyes prior to the time when M1 shifted gaze to look at M2’s eyes (Fig. S8a). Similarly, for *Other Solo Eyes,* we analyzed events when M1’s gaze positions were within 20 visual degrees at any time point after time zero during *Other Solo Eyes* events (Fig. S8b).

At the population level, we calculated the proportion of cells per brain region that showed distinct activity for 1) *Self-follow Mutual Eyes* vs. *Self Solo Eyes* selectively, 2) *Other-follow Mutual Eyes* vs. *Other Solo Eyes* selectively, or 3) both comparisons for at least 5 consecutive bins during the 500 msec after gaze event onset (Fig. 4e; for single-cell PSTHs of *Mutual Eyes*, see Fig. 4d and Fig. S9a). Pair-wise Chi-square tests (FDR-corrected) compared the proportion of all significant cells across brain regions. For each region, we also tested the proportion of significant cells as a function of context in terms of mutual eye contact sequence. For each combination of context and brain region, we further compared the proportion of significant cells that fired more for *Mutual Eyes* to those that fired more for corresponding *Solo Eyes*. The same procedure was repeated for examining activity during a later time period, 500–1000 msec following gaze event onset (Fig. S9b), and for the subset of cells showing ‘social discriminability’ (Fig. S9c).

### Overlap of social gaze interaction functions

To examine the overlap of individual cells involved in the three social gaze interaction variables (social discriminability, social gaze monitoring, and mutual eye contact selectivity; Fig. 4f), we calculated the proportion of cells in each brain region that were involved in none, one, two, or all three functions (Fig. 4g). Within each brain region, we then compared the proportion of cells involved in only one function to at least two functions using Chi-square test with FDR correction. In addition, we compared each of these two proportions of cells across four brain regions.

## Supplemental Materials

**Figure S1.**
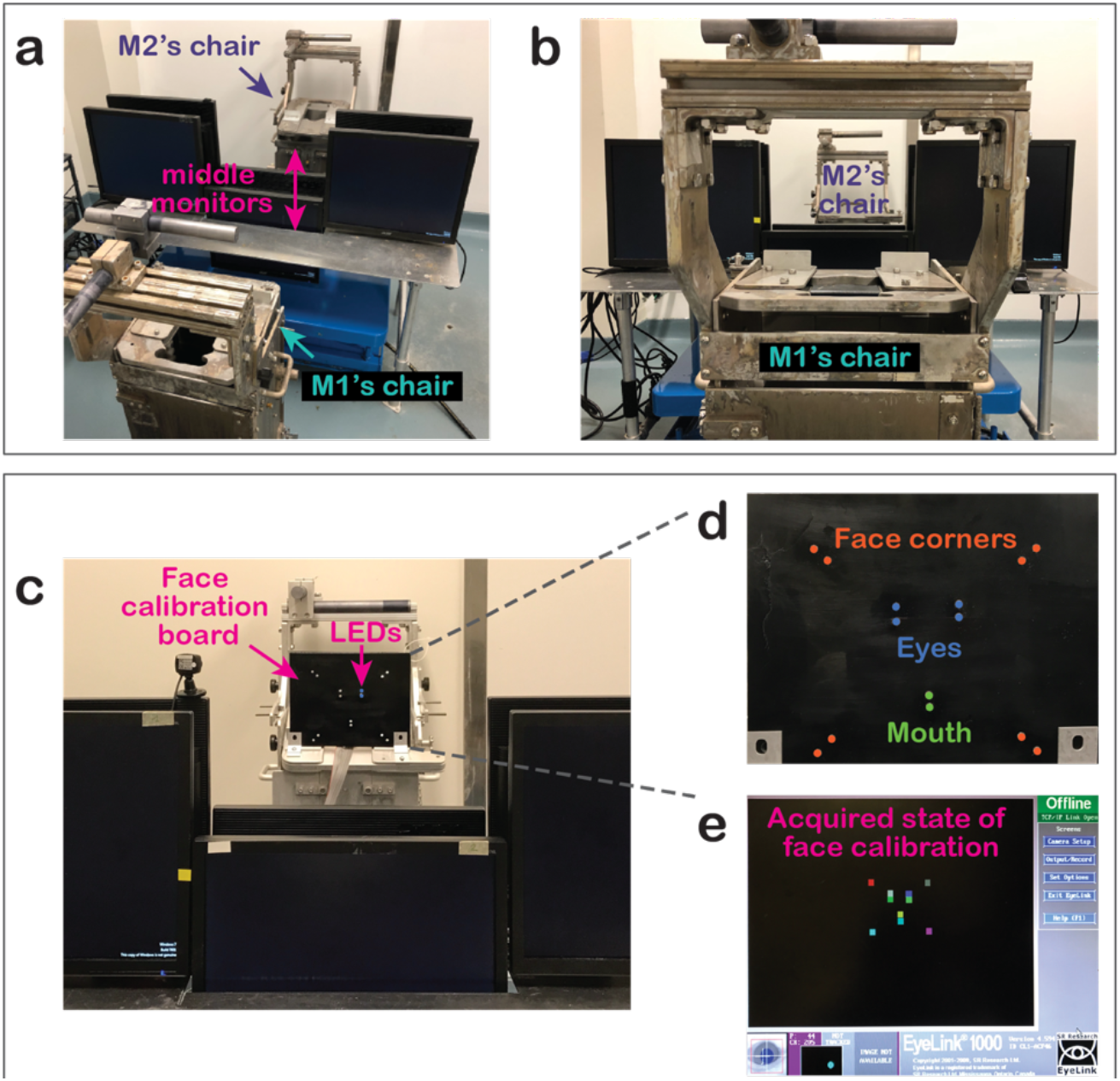
Experimental setup and face calibration board. **a-b**, The experimental setting for live social gaze interaction where a pair of monkeys sat in primate chairs facing each other. The two middle monitors could be lowered down or raised up by using a remote hydraulic system so that the monkeys were able to or not able to see each other. **c–d,** Customized face calibration LED board for face calibration where LED lights were aligned to a monkey’s eyes, mouth, and the four corners of the face. Two sets of LED lights were built to fit two different sizes of monkey faces. **e**, An example state of acquired face calibration, in which each colored square represents an acquired mapping between gaze position and one of the LED positions on the face calibration board.

**Figure S2.**
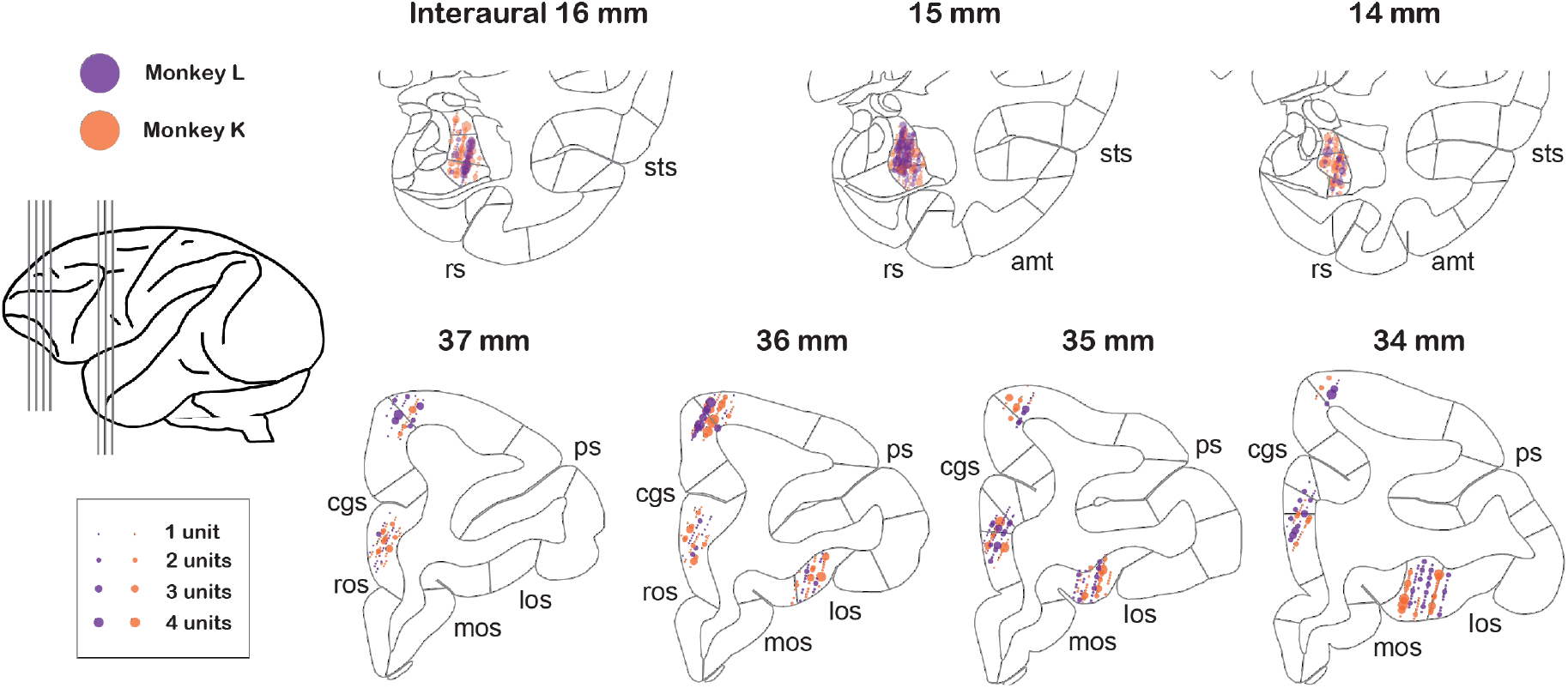
Anatomical locations of recorded single cells in BLA, OFC, ACCg, and dmPFC. Recording locations for individual cells from monkey L (purple points) and monkey K (orange points) projected onto the standard stereotaxic coordinates of the rhesus macaque brain atlas(Paxinos et al., 1999). Three representative coronal slices with 1-mm interaural spacing were chosen for BLA, and four representative coronal slices were chosen for the three prefrontal areas (as shown in the left illustration). Selected landmarks are labeled: cingulate sulcus (cgs), principal sulcus (ps), medial orbitofrontal sulcus (mos), lateral orbitofrontal sulcus (los), superior temporal sulcus (sts), anterior middle temporal sulcus (amt), and rhinal sulcus (rs).

**Figure S3.**
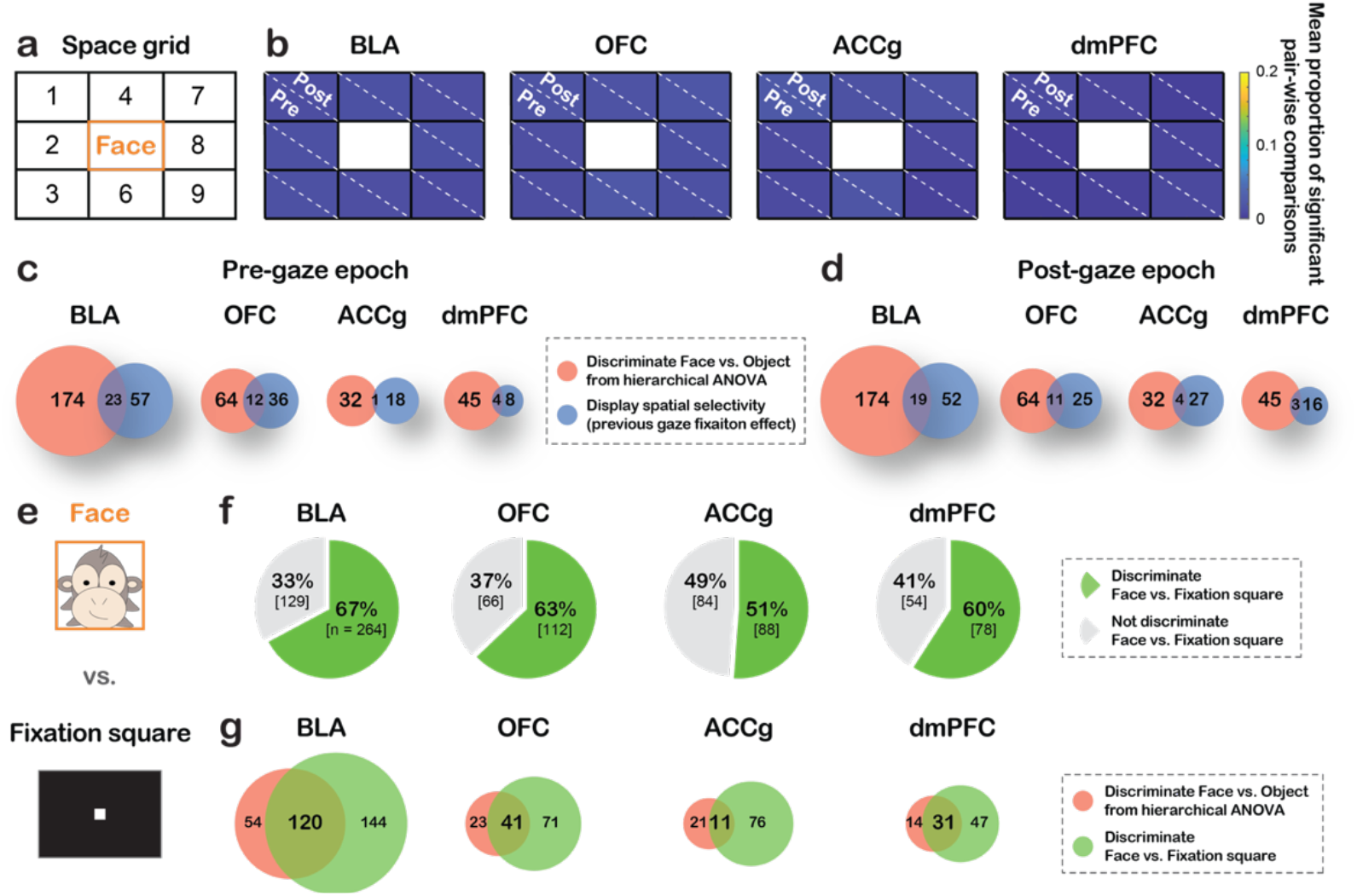
Control analyses for the ROI-based spiking activity analyses. **a–d**, Previous gaze fixation location control, controlling for the location of previous fixation leading up to a current *Face* event. **a**, A 9-cell space grid (labeled as 1–9) was constructed to be centered on partner monkey’s *Face*. We compared neural activity around current *Face* events where the previous fixations fell within different grids. **b**, Heatmaps show the mean proportions of significant pair-wise comparisons among the 8 grids for firing rate differences during the pre-gaze and post-gaze epochs, shown separately split by the diagonal line (e.g., values on the bottom left and top right represent the proportions for the pre-gaze and post-gaze epochs, respectively). None of the 8 grid cells showed any meaningful proportions – across four areas, the mean proportion of 32 cases for pre-gaze epoch was 0.009 ± 0.005 (mean ± std) and that of 32 cases for post-gaze epoch was 0.01 ± 0.004. **c–d,** Venn diagrams show only small overlaps between cells that discriminated *Face* from *Object* based on the hierarchical ANOVA classification (red) and cells that discriminated space grids (blue) for all brain regions separately for the pre-gaze epoch (**c**) (out of cells with ‘social discriminability’ (discriminating *Face* from *Object*): 12% of BLA, 16% of OFC, 3% of ACCg, and 8% of dmPFC) and post-gaze epoch (**d**) (out of cells with ‘social discriminability’: 10% of BLA, 15% of OFC, 11% of ACCg, and 6% of dmPFC), suggesting that activity of cells with ‘social discriminability’ was not simply driven by previous fixation location. **e–g**, Central fixation control. **e,** Neural activity for gazing at partner’s *Face* was compared to looking at a white central *Fixation square* shown on the middle monitor during inter-session breaks (Methods). **f**, Proportions of cells in each of the four brain regions that significantly discriminated *Face* from *Fixation square* (green) and those that did not (gray). Many cells in all four regions differentiated looking at the two stimuli both appearing directly in front of the recorded monkey. **g**, Venn diagrams show some overlaps between cells that discriminated *Face* from *Object* based on the hierarchical ANOVA classification (red) and cells that discriminated *Face* from *Fixation square* (green) for all regions (out of cells with ‘social discriminability’; 69% of BLA, 64% of OFC, 34% of ACCg, and 69% of dmPFC), supporting that activity of cells with social discriminability was unlikely driven by the different visual angle between looking at *Face* and *Object*.

**Figure S4.**
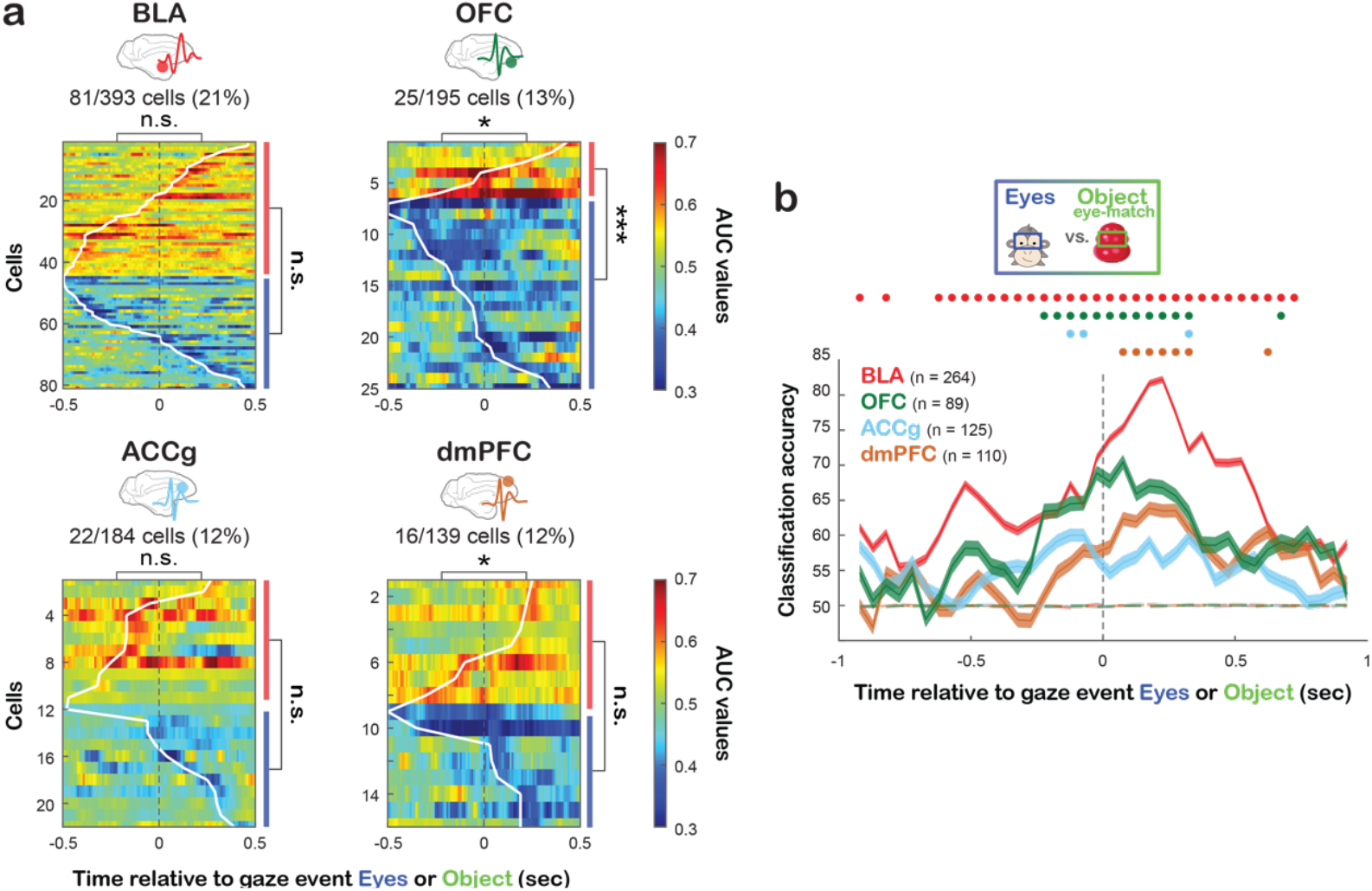
Temporal representations associated with looking at *Eyes* vs. *Object*. **a,** Temporal profiles of spiking activity for *Eyes* vs. *Object* with matching ROI size, same format as Fig. 2a-b. The asterisks on the top of each heatmap indicate the comparisons of the proportions of cells that began discriminating *Eyes* vs. *Object* during the pre-gaze vs. post-gaze epoch. To the right of each heatmap, the red bar represents the proportion of cells with greater activity for *Eyes* and the blue bar for greater activity for *Object* (categorized based on the first-time bin of significant sequence; Methods). *, p < 0.01; ***, p < 0.001; n.s, not significant, Chi-square test, FDR-corrected. **b,** Populations decoding accuracy for *Eyes* vs. *Object* in BLA (red), OFC (green), ACCg (blue), and dmPFC (brown), same format as Fig. 2c–d. Real data are shown in solid lines and empirically derived null data are in dotted lines. Circles at the top indicate the time bins with decoding accuracy significantly higher than the null in corresponding colors (p < 0.001, permutation test).

**Figure S5.**
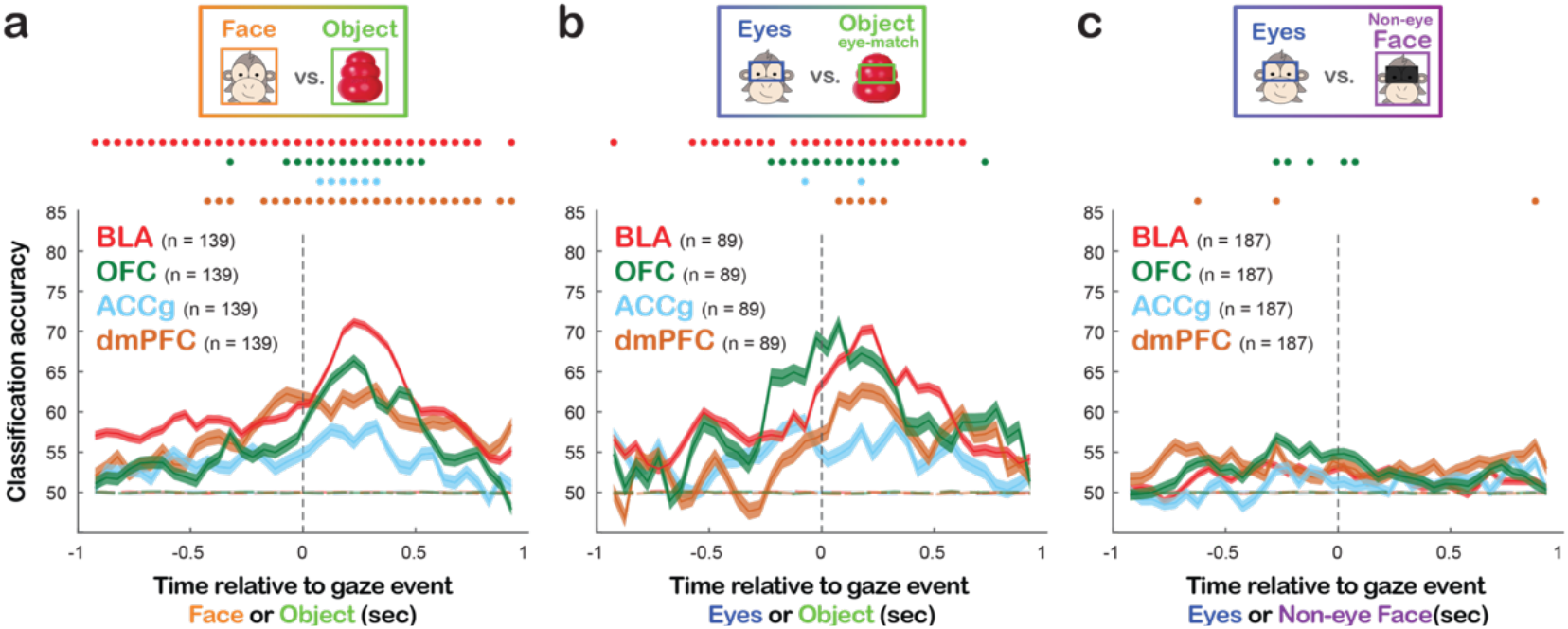
Population decoding accuracy for social gaze events using matching number of cells across regions. **a,** Population decoding accuracy for *Face* vs. *Object* (with matching ROI size) using the matching numbers of BLA (red), OFC (green), ACCg (blue), and dmPFC (brown) cells across the four brain regions (Methods). **b,** Population decoding accuracy for *Eyes* vs. *Object* (with matching ROI size) using the matching number of cells across four regions. Same format as **a. c,** Population decoding accuracy for *Eyes* vs. *Non-eye Face* using the matching number of cells across four regions. Same format as **a.**In all panels, real data are shown using solid lines and empirically derived null data as dotted lines. Circles at the top indicate time bins with decoding accuracy significantly higher than the null in corresponding colors (p < 0.001, permutation test).

**Figure S6.**
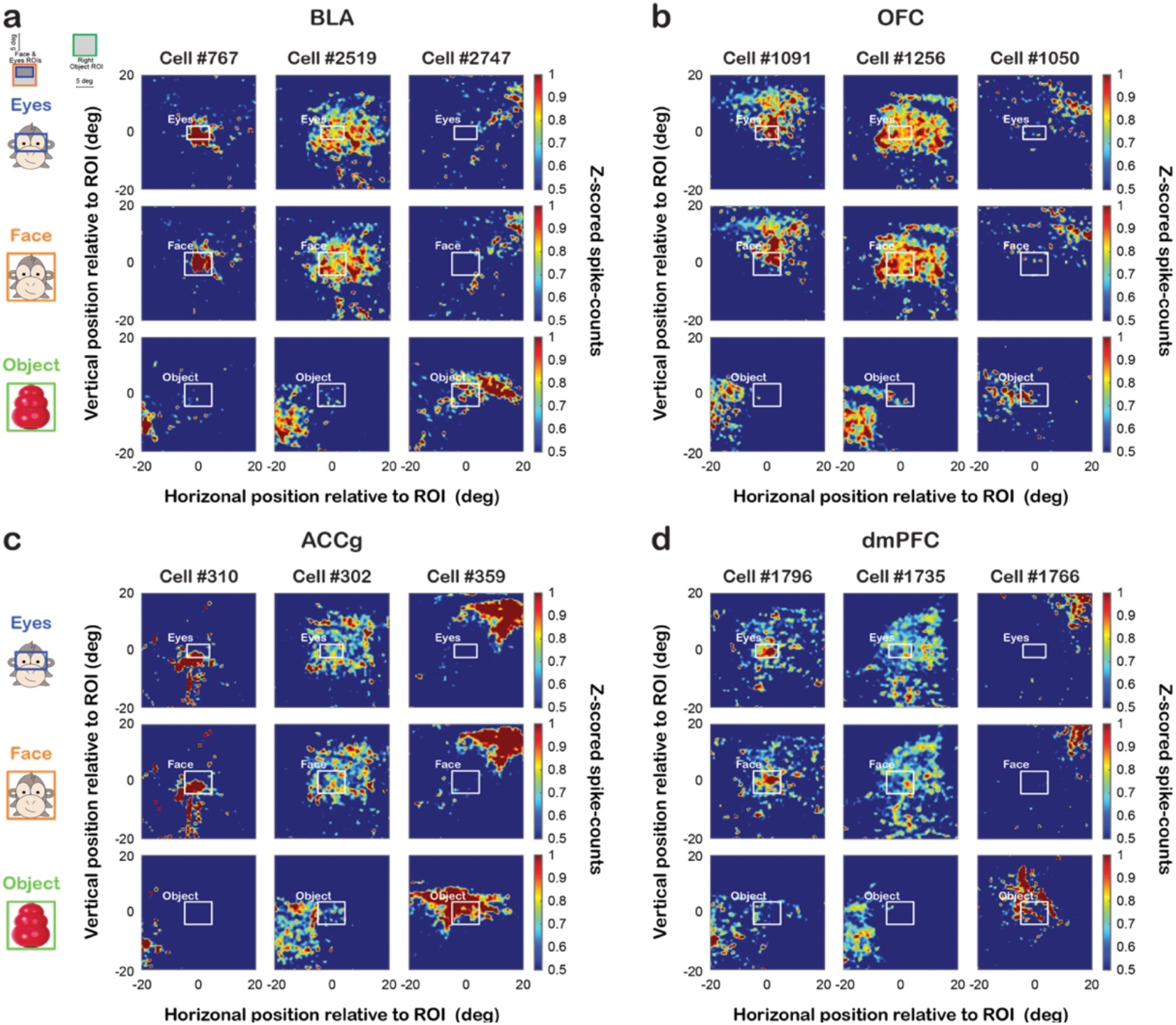
Single-cell examples of spike density maps for *Eyes*, *Face*, and *Object* ROIs. **a-d**, Z-scored spike density heatmaps of three example cells from each brain region, BLA (**a**), OFC (**b**), ACCg (**c**), and dmPFC (**d**), based on recorded monkey’s gaze positions relative to partner monkey’s *Eyes* (first row), *Face* (second row) and *Object* (last row). Top left inset, the layout of the setup for reference. Some cells increased activity selectively when fixating within *Eyes* or *Face*, but not *Object* (first column), while others did so in a more spatially distributed manner (second column). Another group of cells fired more selectively for fixations closer to *Object* (third column).

**Figure S7.**
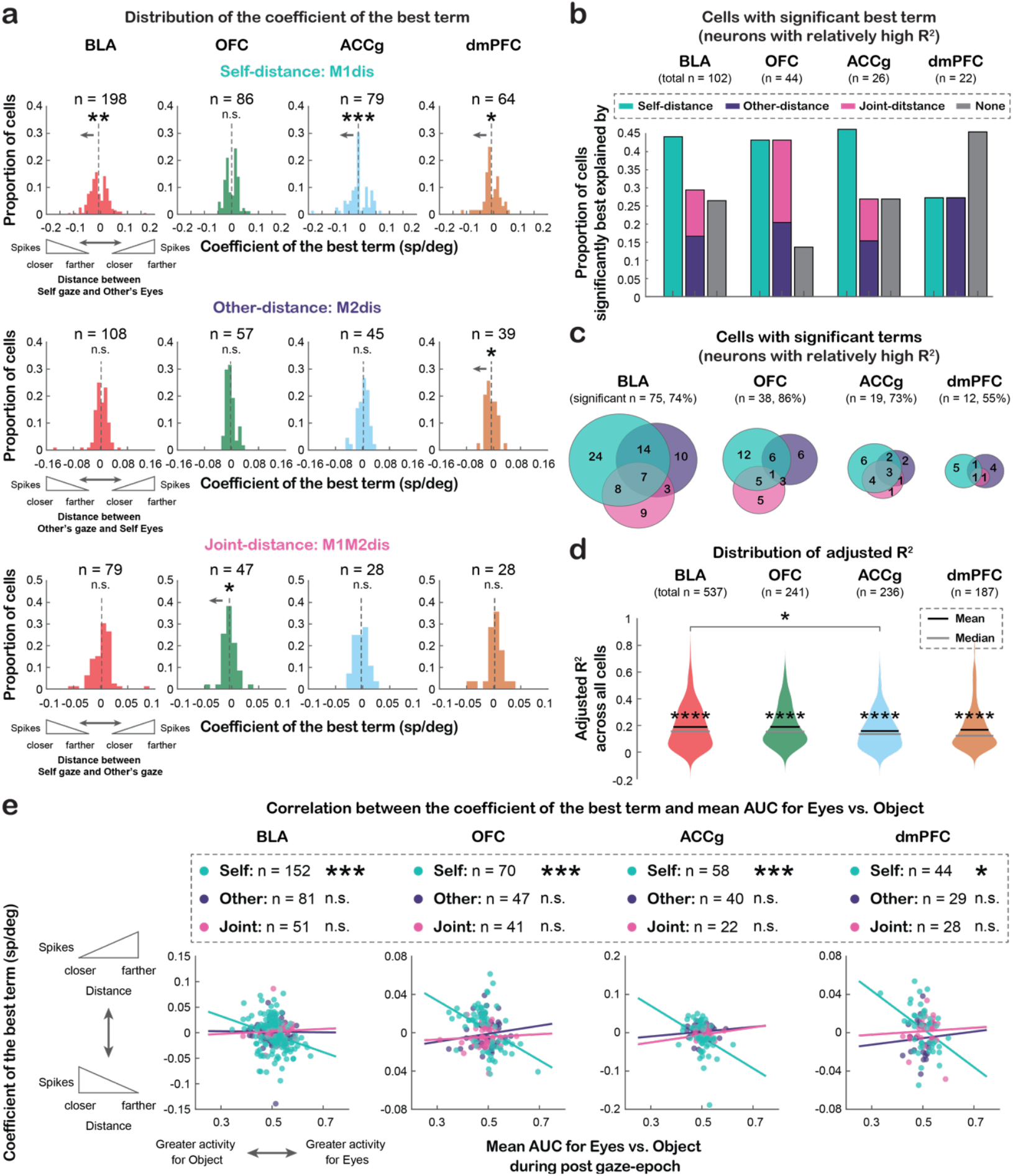
Single-cell correlates of social gaze monitoring from the perspective of self, other, or both (joint). **a**, Distributions of the coefficients of the best significant term when the best term was Self-distance (first row), Other-distance (second row), or Joint-distance (last row) across all cells from each brain region. When considering the best significant term of all cells, BLA, ACCg, and dmPFC showed a population-level bias toward a negative coefficient for Self-distance (t(197) = −2.86, p < 0.005; t(78) = −3.54, p < 0.001; and t(63) = −2.14, p = 0.04, respectively; one-sample t-test, two-sided), indicating that the three populations fired more as the recorded monkey fixated closer to partner’s eyes. In addition, dmPFC showed a bias toward a negative coefficient for Other-distance (t(38) = −2.52, p = 0.02), indicating that dmPFC cells on average had greater activity as the partner monkey looked closer to recorded monkey’s eyes. For Joint-distance, OFC population showed a bias toward a negative coefficient (t(46) = −2.42, p = 0.02), corresponding to higher activity as the gaze positions of self (M1) and other (M2) became closer. *, p < 0.05; **, p < 0.01; ***, p < 0.001, one-sample t-test, two-sided. **b–c**, Same format as Fig. 3d–e, but using a subset of cells with relatively high *R*^*2*^ (adjusted *R*^*2*^ > 0.33) (Methods), resulting in 102 BLA, 44 OFC, 26 ACCg, and 22 dmPFC cells. **b**, Proportions of cells in each region with relatively high *R*^*2*^ whose activity was significantly best explained by Self-distance (mint), Other-distance (purple), Joint-distance (pink), or none (gray). **c**, Venn diagrams of cells with relatively high *R*^*2*^ whose activity could be significantly explained by one or more of the distance variables, regardless of the best-fitting term. **d**, Distributions of adjusted *R*^*2*^ of all cells in each region compared to the null distributions of mean adjusted *R*^*2*^ (always greater, p = 0, permutation test, shown by asterisks ****). We also found a regional difference in the quality of model fits (F(3, 1197) = 3.18, p = 0.02; ANOVA), driven by better fits by BLA than ACCg (p = 0.04, Tukey test). **e**, Correlations between the coefficient of the best term for all cells and the mean AUC values for *Eyes* vs. *Object* during the post-gaze epoch in each brain region. Colored data points represent single cells whose best term was Self-distance, Other-distance, or Joint-distance, with fitted lines from a linear regression shown in corresponding colors.

**Figure S8.**
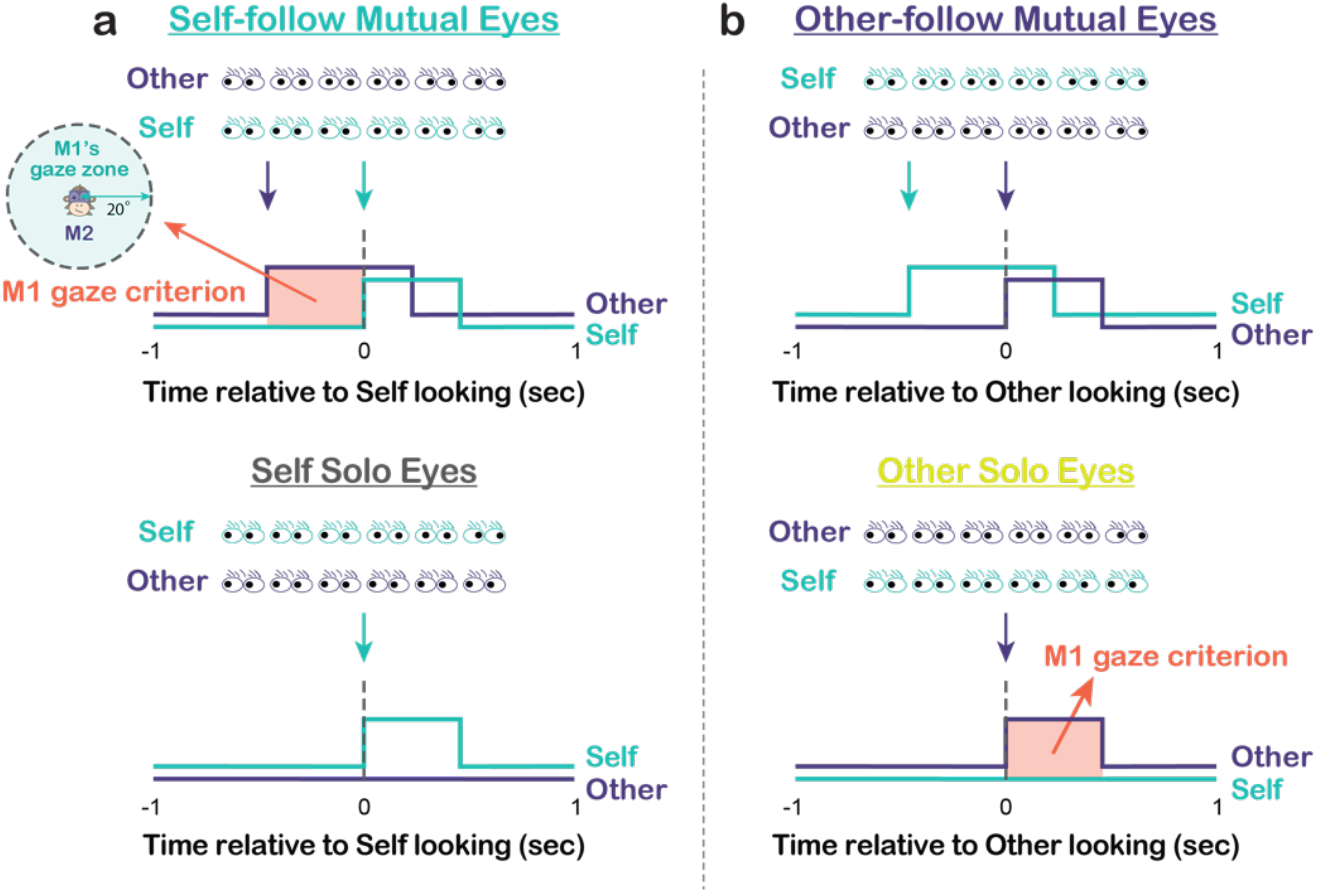
Illustrative diagrams for neural analyses of *Interactive Mutual Eyes* and *Solo Eyes*. **a**, For each cell, spiking activity associated with *Self-follow Mutual Eyes* (mint, top) was compared to *Self Solo Eyes* (gray, bottom), both aligned to the time of M1(self) looking at M2’s (other) eyes. **b**, Similarly, spiking activity associated with *Other-follow Mutual Eyes* (purple, top) was compared to *Other Solo Eyes* (mustard, bottom), both aligned to the time of M2 looking at M1’s eyes. The time segment marked in orange represents the period to which we applied a gaze criterion for M1 (Methods).

**Figure S9.**
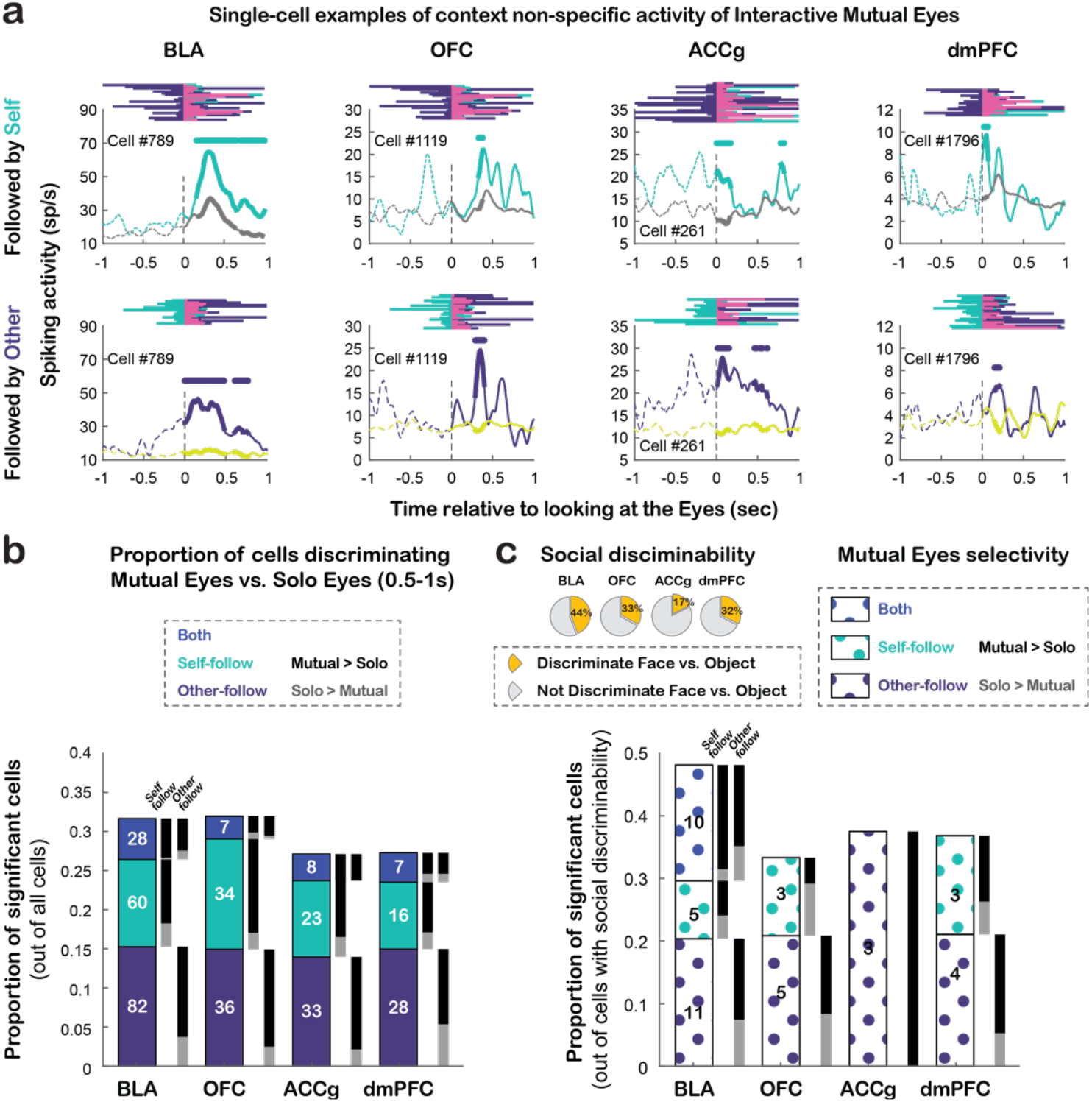
Single-cell examples of context non-specific activity of *Interactive Mutual Eyes* and additional analyses of *Mutual Eyes* selectivity. **a,** Single-cell PSTH examples of context non-specific mutual eye contact selectivity from each of the four brain regions, discriminating both *Self-follow Mutual Eyes* vs. *Self Solo Eyes* and *Other-follow Mutual Eyes* vs. *Other Solo Eyes*. **b**, Proportions of significant cells out of all cells from each area that selectively differentiated *Self-follow Mutual Eyes* from *Self Solo Eyes* (mint), selectively differentiated *Other-follow Mutual Eyes* from *Other Solo Eyes* (purple), or differentiated both types of comparisons (blue, ‘Both’) during 500– 1000 msec following gaze event onset. Same format as Fig. 4e. All four brain regions showed similar proportions of cells for *Self-follow Mutual Eyes* and *Other-follow Mutual Eyes* (all *χ*^2^ < 3.41, p > 0.06, Chi-square test, FDR-corrected). However, there was an effect of agent-specific context in all four regions due to substantially lower proportions of context non-specific *Mutual Eyes* (both types vs. *Self-follow*: *χ*^2^ > *7.26*, p < 0.01 for BLA, OFC, and ACCg, *χ*^2^ *=* 3.52, p = 0.06 for dmPFC; both types vs. *Other-follow*: all *χ*^2^ > 12.60, p < 0.0005, FDR-corrected). As in Fig. 4e, when considering all combinations of agent-specific contexts and brain regions, the majority of significant cells had higher activity for *Interactive Mutual Eyes* than *Solo Eyes* events (all *χ*^2^ > 4.00, p < 0.05 for 11 of 16 cases, FDR-corrected). The black bars indicate the proportions of cells with greater activity for *Interactive Mutual Eyes* than *Solo Eyes*, whereas the gray bars indicate the opposite. **c**, Proportions of significant cells out of all cells with ‘social discriminability’ that selectively differentiated *Self-follow Mutual Eyes* from *Self Solo Eyes*, selectively differentiated *Other-follow Mutual Eyes* from *Other Solo Eyes*, or differentiated both types of comparisons (‘Both’), during the first 500 msec following gaze event onset. Same format as **b**. There were similar proportions of cells that had higher activity for *Interactive Mutual Eyes* and for *Solo Eyes* (all *χ*^2^ < 3.00, p > 0.08 for 8 of 9 cases).

